# Structural basis for recognition of a gonococcal lipooligosaccharide epitope by monoclonal antibody 2C7 informs vaccine and immunotherapeutic design

**DOI:** 10.64898/2026.05.28.728500

**Authors:** Anastasia S. Tsagkarakou, Ferran Nieto-Fabregat, Alessandro Antonio Masi, Yunqin Zhang, Fei Fan, Jutamas Shaughnessy, Peter A. Rice, Xuefei Huang, Sanjay Ram, Peter T. Beernink, Alba Silipo

## Abstract

*Neisseria gonorrhoeae* causes gonorrhea and poses a growing global health threat driven by antimicrobial resistance and the lack of an effective vaccine. The lipooligosaccharide (LOS) epitope recognized by monoclonal antibody (mAb) 2C7 is expressed by most clinical isolates, making it an attractive therapeutic target. To determine the molecular basis for mAb 2C7 recognition, we conducted structural studies of mAb 2C7 in complex with an octasaccharide derived from LOS. NMR epitope mapping demonstrated that mAb 2C7 contacts two of three glycan chains: the β-chain (lactose) and the γ-chain (N-acetylglucosamine). The 1.6-Å resolution crystal structure of the Fab 2C7–octasaccharide complex revealed that these chains make extensive hydrogen bonds and hydrophobic contacts within a cleft of the Fab. Molecular dynamics (MD) simulations showed that the β and γ chains adopt stable, specific conformations, contrasted with transient interactions of the α chain. Informed by these structural data, we synthesized a tetrasaccharide comprising the β and γ chains joined by heptose II. The tetrasaccharide conjugated to a carrier protein bound mAb 2C7 by ELISA and Western blotting. Isothermal titration calorimetry showed that the tetrasaccharide and octasaccharide bound mAb 2C7 with similar affinities, indicating that the smaller oligosaccharide retains the key binding determinants. Binding was enthalpically driven, consistent with the polar interactions observed crystallographically and by MD. NMR experiments with the tetrasaccharide confirmed interactions with all four residues, establishing it as the minimal 2C7 epitope. Together, these studies provide a structural framework for rational design of glycan-based vaccines and antibody therapeutics against antimicrobial-resistant *N. gonorrhoeae*.

## INTRODUCTION

*Neisseria gonorrhoeae* (the gonococcus) causes gonorrhea, the second most prevalent bacterial sexually transmitted infection with approximately 80 million cases worldwide annually [1, 2]. Gonococcal infections cause urethritis in men, cervicitis and pelvic inflammatory disease in women and, infrequently, disseminated infections [3]. A major challenge is the high incidence of asymptomatic infections in the rectum and pharynx, and the female genital tract, which cause unrecognized transmission and sustained disease burden at the population level. Gonococcal strains have developed resistance to all major classes of antibiotics, including β-lactams, fluoroquinolones, macrolides and cephalosporins, which raise the threat of untreatable gonococcal infections [3-5]. The progressive erosion of treatment options underscores the need for effective vaccines and novel therapeutic strategies, including non-antibiotic approaches.

The search for conserved gonococcal vaccine targets is challenging because many surface-exposed proteins undergo antigenic variation. However, several conserved surface-exposed proteins have been shown to elicit bactericidal antibodies or reduce bacterial burden in experimental models. These proteins include surface-exposed lysozyme inhibitor of C-type lysozyme (SliC) conjugated to virus-like particles [6], methionine-binding protein (MetQ) [7], Neisserial heparin-binding antigen (NHBA) [8], multidrug efflux transporter (E subunit) MtrE [9], macrophage infectivity potentiator (MIP) [10], and adhesin complex protein (ACP) [11]. Outer membrane vesicles (OMV) containing surface proteins from either gonococci or meningococci have been investigated in preclinical studies to evaluate protection against gonococcal diseases [12-14] and several OMV vaccines have advanced to clinical trials [15].

Lipooligosaccharide (LOS) is the most abundant molecule in the outer membrane of gonococci and is an important virulence factor [16]. Gonococcal LOS consists of a conserved lipid A core and three variable glycan chain extensions. The lipid A component induces intense inflammatory responses by stimulating the Toll-like receptor 4 (TLR4)/MD2 receptor complex. The core region includes two 2-keto-3-deoxy-D-manno-octulosonic acid (KDO) and two heptose residues, HepI and II (**Fig. S1**). The two Hep residues bear three glycan chain extensions. The alpha (α) chain extends from HepI and comprises two to five additional monosaccharides. The beta (β) chain extends from the 3-position of HepII and consists of zero, two or (in rare instances) four [17] additional sugars. The gamma (γ) chain extends from the 2-position of HepII and comprises a GlcNAc residue.

The oligosaccharide (OS) component is antigenically variable due to phase variation of four LOS glycosyltransferase genes (*lgtA, lgtC, lgtD and lgtG)*, which results in different glycan extensions among strains and sometimes within the same strain [18, 19]. In addition, gonococci scavenge cytidine monophospho (CMP)-sialic acid from the human host during infection, which they use to sialylate terminal Gal residues of LOS [16]. Sialylation of the α chain lacto-N-neotetraose extension enhances complement Factor H (FH) binding, which down-regulates amplification of the alternative pathway leading to increased resistance of the bacteria to complement-mediated killing [20, 21]. Sialylation of the β chain lactose extension limits complement C3 deposition and enhances binding to Siglec receptors on host cells [22]. Thus, both genetic and host-derived modification of LOS enhances antigenic variability and immune evasion.

Anti-LOS mAbs that recognize different LOS structures have been described [16]. One of these mAbs, known as 2C7 [23] binds to LOS that bears a β-chain lactose extension [24]. mAb 2C7 mediates complement-dependent bactericidal activity against almost all minimally passaged clinical isolates [22, 25] and decreases bacterial burden in the mouse vaginal colonization model [16, 26]. A human IgG1 chimeric derivative of mAb 2C7 decreased gonococcal vaginal colonization in mice [27]. To guide humanization of mAb 2C7, we determined NMR and crystal structures of chimeric mAb (or Fab) 2C7 alone or bound to a peptide mimic of the 2C7 LOS epitope [28]. These structures provided detailed information on binding of the peptide mimic, however an understanding of the molecular basis for recognition of the carbohydrate epitope recognized by mAb 2C7 remained elusive. Here, we report comprehensive binding and structural analyses of mAb 2C7 (or its antigen-binding fragment, Fab) and two oligosaccharides (OS): an octasaccharide derived from acid hydrolysis of LOS and a synthetic tetrasaccharide.

## RESULTS

### Purification and characterization of *N. gonorrhoeae* OS

The LOS of *N. gonorrhoeae* strain 15253 was extracted and purified as described in the Methods (Section 2.2) [29]. SDS-PAGE followed by silver nitrate staining [30] confirmed the rough nature of the extracted LPS (R-LPS or LOS), as indicated by the low-molecular-weight band observed at the bottom of the gel (**Fig. S4**). Compositional analysis [31] revealed the presence of Galactose (Gal), Glucose (Glc) N-Acetylglucosamine (GlcNAc), heptose (Hep), and 3-deoxy-D-manno-oct-2-ulosonic acid (KDO). LOS was subjected to mild acid hydrolysis to selectively cleave the acid-labile glycosidic bond between the KDO and the non-reducing GlcN of the lipid A region. Following several purification steps, the OS fraction was analyzed by negative-mode MALDI-TOF MS (**Fig. S5**). The MS analysis indicated that the main species (*m/z* 1577.53) is an octasaccharide containing 1 KDO, 2 Hep, 4 Hex, 1 HexNAc, and 1 phosphoethanolamine (PEtN), consistent with a previous report [31]. NMR spectra of the purified octasaccharide showed the presence of anomeric signals (**Table S2** and **Fig. S6**) attributable to seven different spin systems, in agreement with the composition determined via MS analysis (**Fig. S5**).

### NMR reveals the glycan epitope recognized by mAb 2C7

The molecular recognition of purified gonococcal octasaccharide by mAb 2C7 was unveiled using a combination of NMR experiments and MD simulations. We employed ligand-based NMR approaches [32, 33] to investigate the molecular details of the interaction. These NMR approaches identified the glycan residues that comprised the binding interface with mAb 2C7 to map the octasaccharide epitope. STD NMR and tr-NOESY provided information on the epitope recognized by the mAb and the conformation of the glycan ligand, respectively. STD NMR experiments revealed that recognition occurred primarily through the β chain of the octasaccharide (**Fig. 1**). The strongest STD signals were attributed to H2, H3, and H4 protons of the terminal βGal residue in the β chain, followed by moderate contributions from H5 and minor involvement from the remaining ring protons. The αGlc at the reducing end of the β chain also showed STD signals, particularly at H2, H3, and H4, indicating a partial contribution to the binding event. The γ chain, αGlcNAc, exhibited the second strongest STD signals, with a prominent effect at H2 and smaller contributions at H3 and H4. In contrast, the α chain exhibited weak or unresolvable STD signals, likely due to spectral overlap and low STD intensities (<10%), which suggested that it did not contribute significantly to the binding interaction. Overall, these data showed that mAb 2C7 primarily recognized the βGal residue of the β chain and the αGlcNAc of the γ chain.

**Figure 1.**
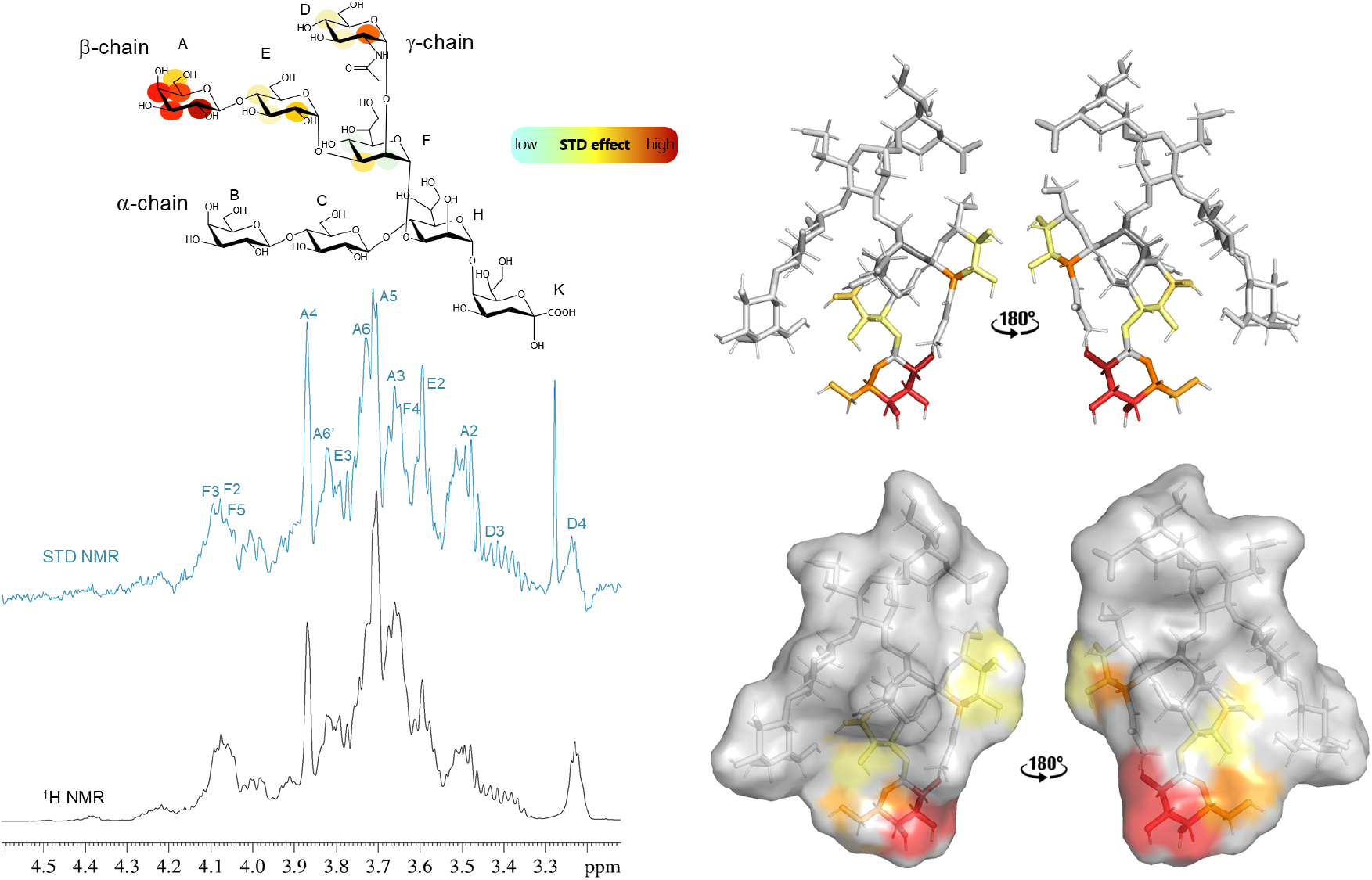
STD NMR analysis and epitope mapping of gonococcal octasaccharide. Top left, schematic view of octasaccharide colored according to STD values. Bottom left, ^1^H and STD spectra. %STD are obtained by the ratio (I_0_ − I_sat_)/I_0_, where (I_0_ − I_sat_) is the signal intensity in the STD-NMR spectrum (blue) and I_0_ is the peak intensity of the off-resonance spectrum (black). Top right, representative 3D structure of the glycan ligand in stick rendering colored according to STD signal intensity. Bottom right, same glycan ligand in surface rendering.

### Crystal structure of Fab 2C7-octasaccharide complex

To identify the atomic interactions of the mAb 2C7 with the LOS glycan and to elucidate the paratope, we crystallized Fab 2C7 in a complex with the octasaccharide and determined its X-ray structure at 1.6-Å resolution by molecular replacement. The crystals are in space group C2 and there are two copies of the Fab-octasaccharide complex in the asymmetric unit. We refined the structure to a working R factor of 19.3% and a free R factor of 21.0%. The data collection and refinement statistics are given in **Table S1**. After cleavage of the signal sequences, the heavy chain (HC) contains residues 25-245 and the light chain (LC) contains residues 21-235. HC loop residues 154-166 and 209-222 were absent from chain A, carboxyl-terminal residues 241-245 were absent from chain A and residue 245 was absent from chain C. LC carboxyl-terminal residues 232-235 were absent from chain B and terminal residues 1 and 235 were absent from chain D.

A global view of the complex shows that the octasaccharide is bound predominantly to the Fab HC (**Fig. 2A**). The three complementarity determining regions (CDR) of the HC mediate polar and van der Waals interactions with the octasaccharide and contribute an average of 501 Å^2^ of buried surface area to the interaction. The CDR3 loops of the LC mediate additional van der Waals interactions and provide an average of 55 Å^2^ of buried surface area. The contributions of each monosaccharide to the interaction surface area and to polar interactions in the Fab-octasaccharide complex are given in **Table S3**.

**Figure 2.**
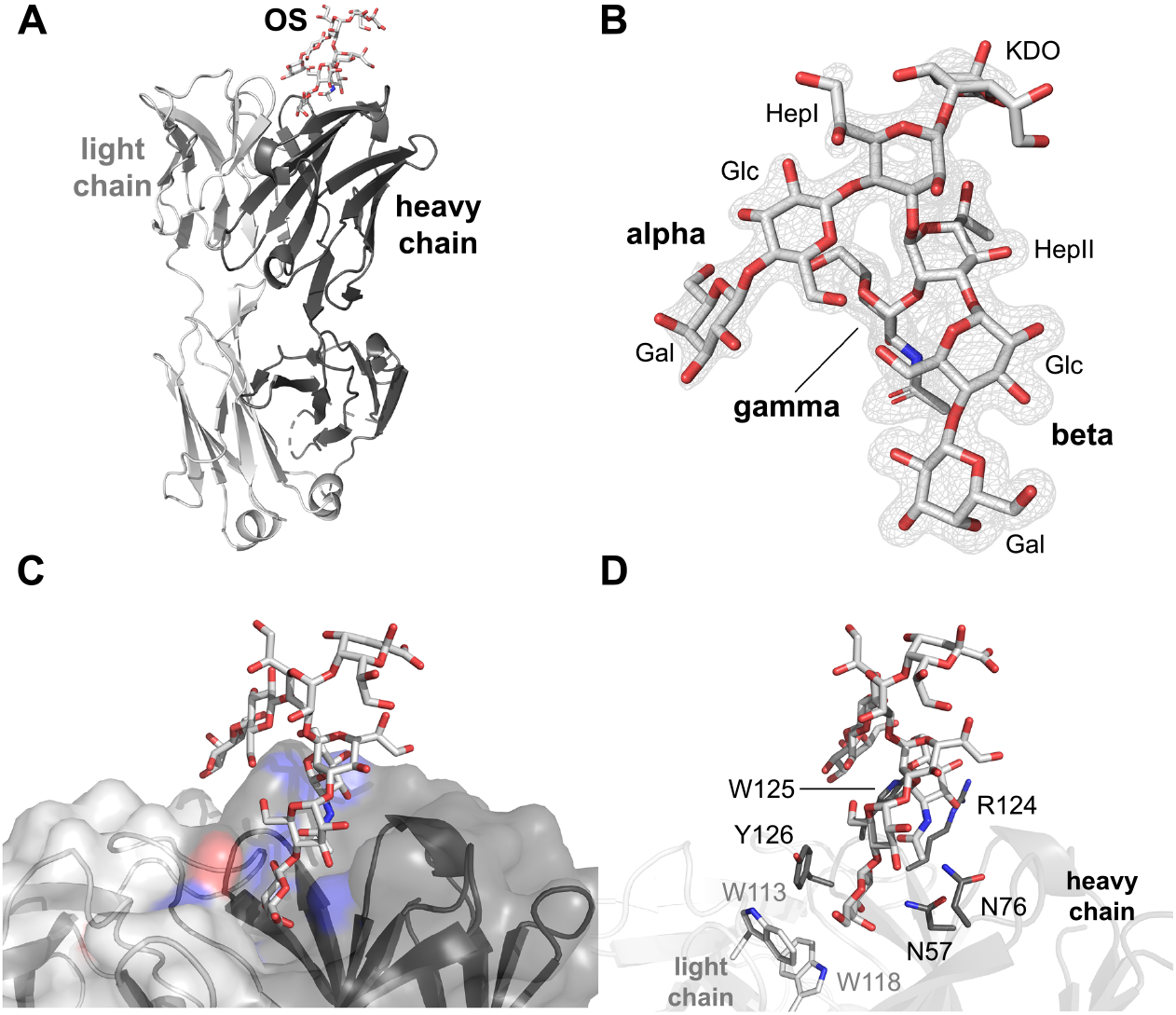
Crystal structure of gonococcal octasaccharide bound to Fab 2C7. **A**, Overall structural view. The Fab heavy chain (HC) is shown in dark grey, the light chain (LC) is shown in light grey and the octasaccharide is colored by atom. The Fab interacts with the octasaccharide primarily via the HC. **B**, Shake-omit electron density map (mFo-DFc) calculated after removing glycan atoms from model (contour level 3 sigma). The abbreviations for the sugar residues are shown in regular typeface and the names of the heptose-linked chains are denoted in bold. **C**, View of the octasaccharide and a surface representation of the Fab. The β and γ chains protrude into a cleft of the Fab. **D**, View of the octasaccharide in relation to Fab side chains involved in polar (N57, N76, R124) or hydrophobic interactions (W113, W118 (LC) and W125, Y126 (HC). The orientations of the octasaccharide in the four panels have been rotated slightly around the Y-axis for clarity.

The electron density for the octasaccharide is well defined as seen in a shake-omit mFo-DFc electron density map (**Fig. 2B**). The KDO residue at the reducing end is not in contact with the Fab and exhibits weaker density than the rest of the octasaccharide. The non-reducing ends are oriented with the β chain (lactose) and γ chain (αGlcNAc) buried in the cleft formed by the complementarity determining region (CDR) loops of the Fab (**Fig. 2C**). The paratope comprises polar (N57, N76 and R124) and aromatic (W125 and Y126) residues from the HC, as well as aromatic (W113 and W118) residues from the LC (**Fig. 2D**).

The galactose (βGal) residue at the non-reducing end of the α chain stacks against W125 of the HC (**Fig. 3A**). Similarly, the βGal residue of the β chain stacks against Y126 of the HC (**Fig. 3B**). Five H bonds are formed between the β chain and the Fab HC, including the side chains of residues N57, N76 and the backbone amide group of Y126 (**Fig. 3C** and **Table S4**). Four H bonds are made between the γ chain and the side chains of R124 and W125 (**Fig. 3D** and **Table S4**). In addition, there are nine water-mediated H bonds between the octasaccharide and the Fab, which involve seven HC residues (**Table S5**). All the H bonds, direct and water mediated, are present in both copies in the asymmetric unit. The glycan also forms 66 van der Waals interactions with side chains atoms of HC residues D55, N57, E59, V74, N76, S123, R124, W125 and Y126.

**Figure 3.**
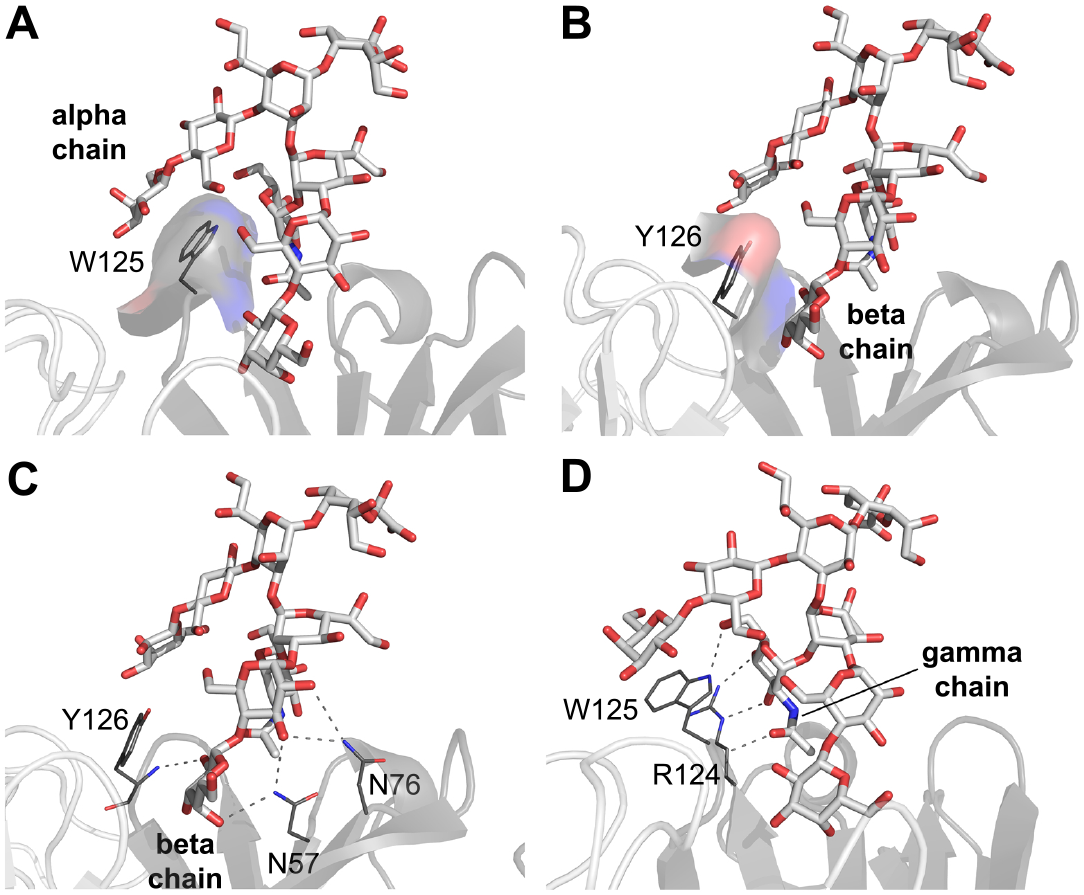
Hydrophobic and H-bond interactions in Fab 2C7-octasaccharide complex. **A**, Octasaccharide α chain terminal βGal residue stacks against residue W125 of the Fab heavy chain (HC; dark gray ribbons). **B**, β chain terminal βGal residue stacks against residue Y126 of the Fab HC. **C**, Fab HC residues N57, N76 and Y126 mediate H bonds with β chain. **D**, Fab HC residues R124 and W125 mediate H bonds with γ chain. The orientations of the octasaccharide in each panel have been rotated slightly around the Y-axis for clarity.

### Molecular dynamics simulations identify glycan conformational variability

We conducted tr-NOESY NMR and computational studies to investigate ligand conformation and binding mode, and to propose a 3D model of the complex combining the spectroscopic data with prior crystallographic data. Comparison of NOE contacts in the free and the bound states revealed no significant differences in interproton distances, indicating that the octasaccharide does not undergo substantial conformational changes upon binding to mAb 2C7. Based on the NMR data, the conformation of the octasaccharide was first studied in the free state. The octasaccharide was built according to the energetic minima calculated through the adiabatic maps by molecular mechanics calculation and subjected to a molecular dynamics (MD) simulation to explore its conformational behavior in solution. To model the complex, the crystallographic structure of Fab 2C7 alone (PDB 8DOZ) [28] was used as the starting protein structure (**Fig. S7**). The octasaccharide model derived from NMR and molecular mechanics calculations was positioned in the Fab binding site and the resulting complex was optimized using the Maestro program suite. The optimized complex was then subjected to MD simulations (500 ns) to refine the interaction and evaluate the stability of the complex based on experimental data. During the MD simulation, *inter*-residue contacts, glycosidic ϕ/ψ torsion angles, and root-mean-square deviation (RMSD) values were sampled to provide insights into the bioactive conformational behavior of the octasaccharide when it interacts with Fab 2C7. RMSD analysis performed on both protein backbone and glycan residues, using the first frame as a reference and evaluating both the octasaccharide and its heptose extensions (a, β and γ chains), indicated a clear stability within the binding pocket of all the segments. The simulated average inter-residue distances, calculated using the ⟨r ^-6^⟩ values, showed a good correlation with the acquired experimental data. The energetically accessible conformational spaces around the glycosidic linkages were explored through molecular mechanics (MM) calculations. The conformational space of the glycosidic linkages was first explored by molecular mechanics calculations, which identified the energetically accessible minima corresponding to conformations stabilized by the *exo*-anomeric effect (**Fig. S8**). Analysis of the glycosidic φ/ψ torsions sampled during the MD simulations revealed that the ϕ angles predominantly adopted values within these low-energy regions, in agreement with the *exo*-anomeric conformation inferred from the experimentally observed inter-residue ROE contacts (**Fig. 4** and **Table S6**).

**Figure 4.**
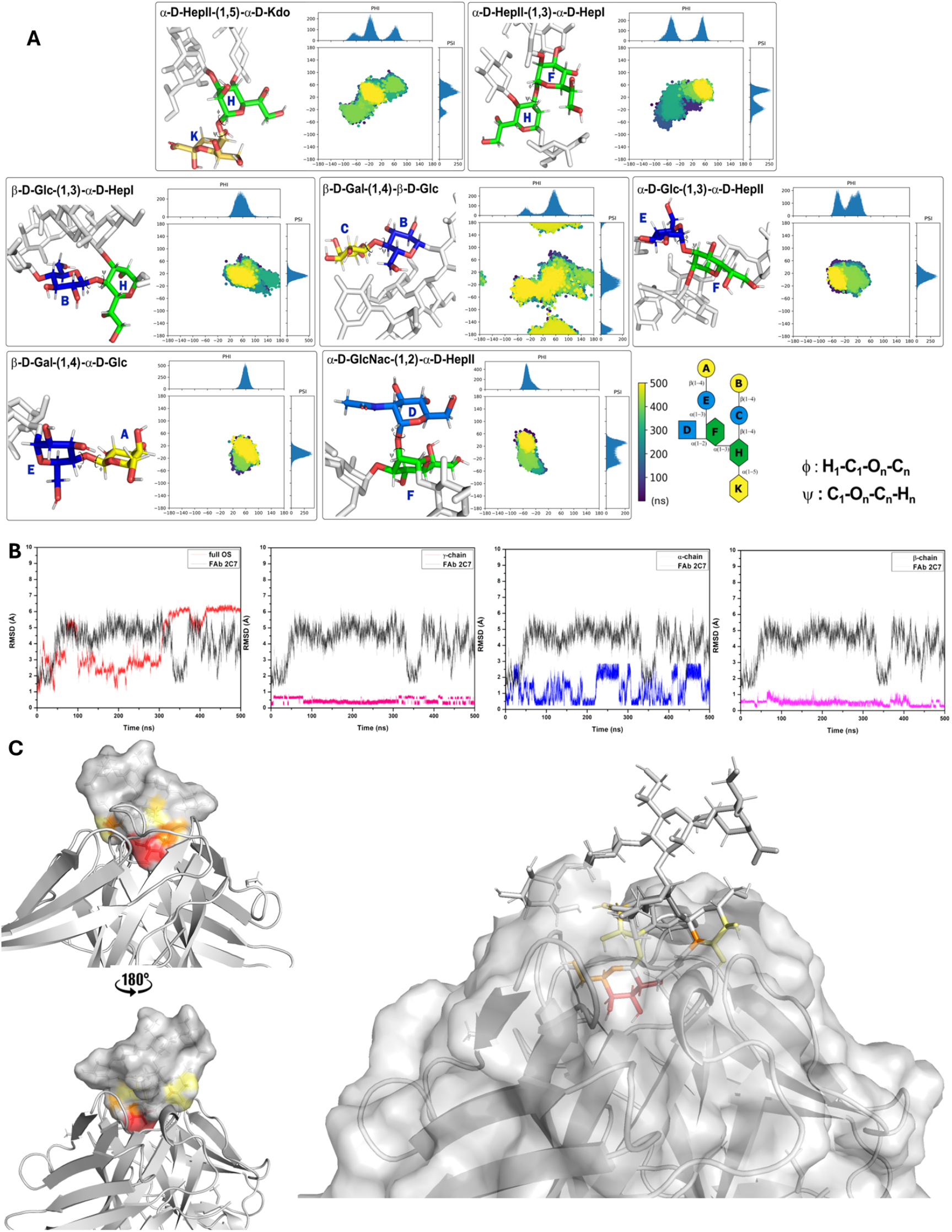
**A**, Dihedral angles analysis represented as scatter plots of the φ torsion against ψ, for the 500 ns MD simulation with the relative histograms representing the most populated regions. **B**, Root mean square deviation (RMSD) plots from molecular dynamics simulations for each chain, depicting the stability and conformational fluctuations of both the ligand and the antibody during the simulation timeframe (500 ns). The antibody data are the same in all four subpanels. **C**, trROESY-derived bioactive conformation of octasaccharide colored according to the STD effects in complex with Fab 2C7.

Cluster analysis based on ligand RMSD was performed on the MD trajectory, and a representative structure of the most populated cluster was selected to illustrate the 3D complex **(Fig. S9A, B)**. Backbone superpositions of a representative pose from each of the most populated clusters showed highly similar interactions between the octasaccharide and the Fab (**Fig. S9C**), indicating good convergence of the simulations. The NOE-based conformation of OS was preserved within the 2C7 binding site during MD simulation. Analysis of the interactions revealed that mAb 2C7 primarily recognized the βGlc and αGal residues of the β-chain, and the αGlcNAc of the γ-chain through polar and hydrophobic interactions. The α-chain was largely solvent exposed, and the KDO residue did not form any interactions with the Fab. However, occasional transient contacts between α-chain terminal βGal residue and Trp125 (∼5% of the trajectory) were observed during the MD simulation. Analysis of H bonds revealed several persistent interactions involving the β-chain; for instance, the hydroxyl group at position 3 of αGlc interacted with Asn56 for the 49% of the trajectory frames. Further, predominant H bonds were mainly associated with the terminal βGal residue of the β-chain. Strong interactions sampled during MD simulation were observed between βGal-H4 and Glu59 (58%), βGal-O2 and Tyr126 (49%), βGal-O4 and Asn57 (45%) and βGal-H2 and Ser123 (31%). Moreover, a strong interaction was found between αGlcNAc-O3 and Asp55 (85%) as well as interactions involving αGlcNAc-O6 and Trp125. The reported percentages correspond to the fraction of simulation frames in which the corresponding H bond was detected. Most interactions involved residues located in the HC variable region (VH) of Fab 2C7, except for a single interaction involving the side chain of Trp118 in the LC.

### Comparison of crystal structure and molecular dynamics ensemble

We observed a close agreement between the crystal structure and a representative pose from the MD ensemble (**Fig. S10**). An extensive network of polar and non-polar interactions was found between the octasaccharide, especially the β and γ chains, and the cleft of the Fab. Most of the polar contacts was found in both structures (**Table 1**). The paratope contains aromatic residues, particularly Trp125 and Tyr126, which provide a hydrophobic scaffold for the binding of sugar rings from the α and β chains. The complementary structural approaches confirmed the STD-derived binding epitope.

**Table 1.**
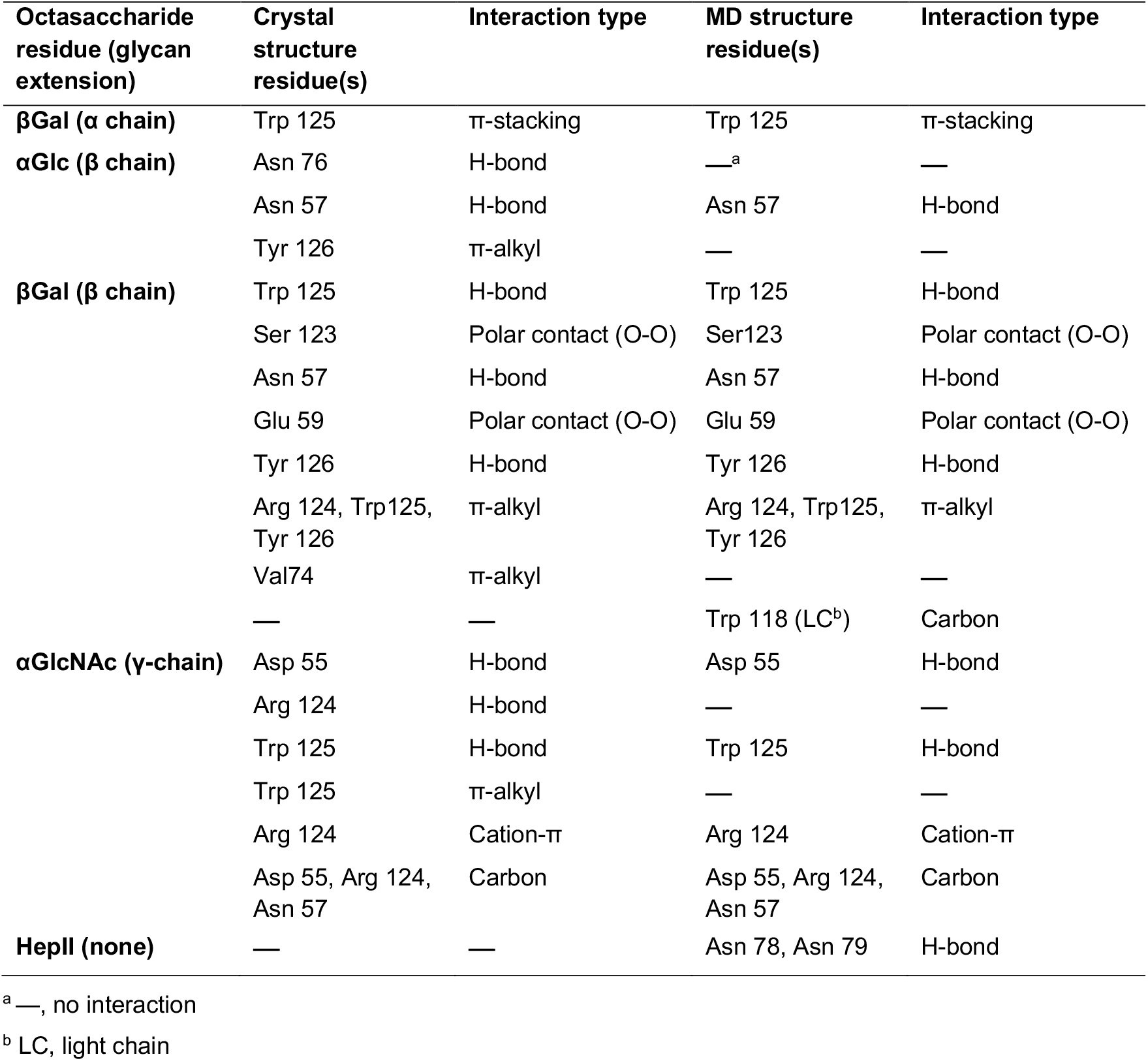
Comparison of interactions between Fab and octasaccharide between crystal and MD structures.

| Octasaccharide residue (glycan extension) | Crystal structure residue(s) | Interaction type | MD structure residue(s) | Interaction type |
| --- | --- | --- | --- | --- |
| βGal ($\alpha$ chain) | Trp 125 | π-stacking | Trp 125 | π-stacking |
| αGlc ($\beta$ chain) | Asn 76 | H-bond | —<sup>a</sup> | — |
|  | Asn 57 | H-bond | Asn 57 | H-bond |
| | Tyr 126 | π-alkyl | — | — |
| βGal ($\beta$ chain) | Trp 125 | H-bond | Trp 125 | H-bond |
|  | Ser 123 | Polar contact (O-O) | Ser123 | Polar contact (O-O) |
|  | Asn 57 | H-bond | Asn 57 | H-bond |
|  | Glu 59 | Polar contact (O-O) | Glu 59 | Polar contact (O-O) |
|  | Tyr 126 | H-bond | Tyr 126 | H-bond |
| | Arg 124, Trp125, Tyr 126 | π-alkyl | Arg 124, Trp125, Tyr 126 | π-alkyl |
| | Val74 | π-alkyl | — | — |
|  | — | — | Trp 118 (LC<sup>b</sup>) | Carbon |
| αGlcNAc (γ-chain) | Asp 55 | H-bond | Asp 55 | H-bond |
|  | Arg 124 | H-bond | — | — |
|  | Trp 125 | H-bond | Trp 125 | H-bond |
| | Trp 125 | π-alkyl | — | — |
| | Arg 124 | Cation-π | Arg 124 | Cation-π |
|  | Asp 55, Arg 124, Asn 57 | Carbon | Asp 55, Arg 124, Asn 57 | Carbon |
| HepII (none) | — | — | Asn 78, Asn 79 | H-bond |

<sup>a</sup> —, no interaction
<sup>b</sup> LC, light chain

The α chain is partially exposed to solvent in both structures and is dynamic along the MD simulation. This chain forms water-mediated H bonds and π-alkyl interactions with the side chain atoms of Trp125. The *β* chain is buried in the cleft of the Fab and its αGlc residue forms H bonds with the side chain atoms of Asn57 and Asn76, and a π-alkyl interaction with the side chain atoms of Trp125. The *β*Gal residue forms H bonding interactions with the side chain atoms of Asn57, Glu59, Ser123 and Tyr126, and forms *π*-alkyl and *π*-*π* stacking interactions with Arg124, Trp125, Tyr126 and Val74. The γ chain is also buried in the cleft of the Fab and its αGlcNAc forms H-bond interactions with the side chain atoms of Asp55, Arg124 and Trp125. The γ chain is further stabilized by extensive π-alkyl contacts with the indole ring of Trp125. Overall, binding studies of the octasaccharide with mAb 2C7 and MD simulations with Fab 2C7 confirmed the binding epitope and the bioactive conformation as described above.

### mAb 2C7 binds a synthetic tetrasaccharide

The NMR data and the crystal and MD structures showed extensive interactions of mAb (or Fab) 2C7 with the β and γ chains (**Tables 1, S4** and **S5**). To determine whether these chains comprised the minimal epitope for binding of mAb 2C7, we synthesized a tetrasaccharide that contains HepII linked to β (lactose) and γ (GlcNAc) chains (**Fig. S2**). This tetrasaccharide was functionalized with an N-hydroxysuccinimide (NHS) linker, which was subsequently conjugated with BSA. An average of 16 tetrasaccharides per BSA was calculated from MALDI-TOF MS analysis (**Fig. S3A**). mAb 2C7 bound to the tetrasaccharide-BSA conjugate in a concentration-dependent manner by ELISA (**Fig. 5A**). mAb 2C7 also bound to the conjugate by Western blotting (**Fig. 5B**), which yielded a size (∼80 kDa) consistent with that from MALDI-TOF analysis (**Fig. S3A**). Smaller amounts of higher molecular mass oligomers were also present. Collectively, these results were consistent with the NMR studies that showed STD signals for the β and γ chains and confirmed that the tetrasaccharide was likely to be the minimal 2C7-binding epitope.

**Figure 5.**
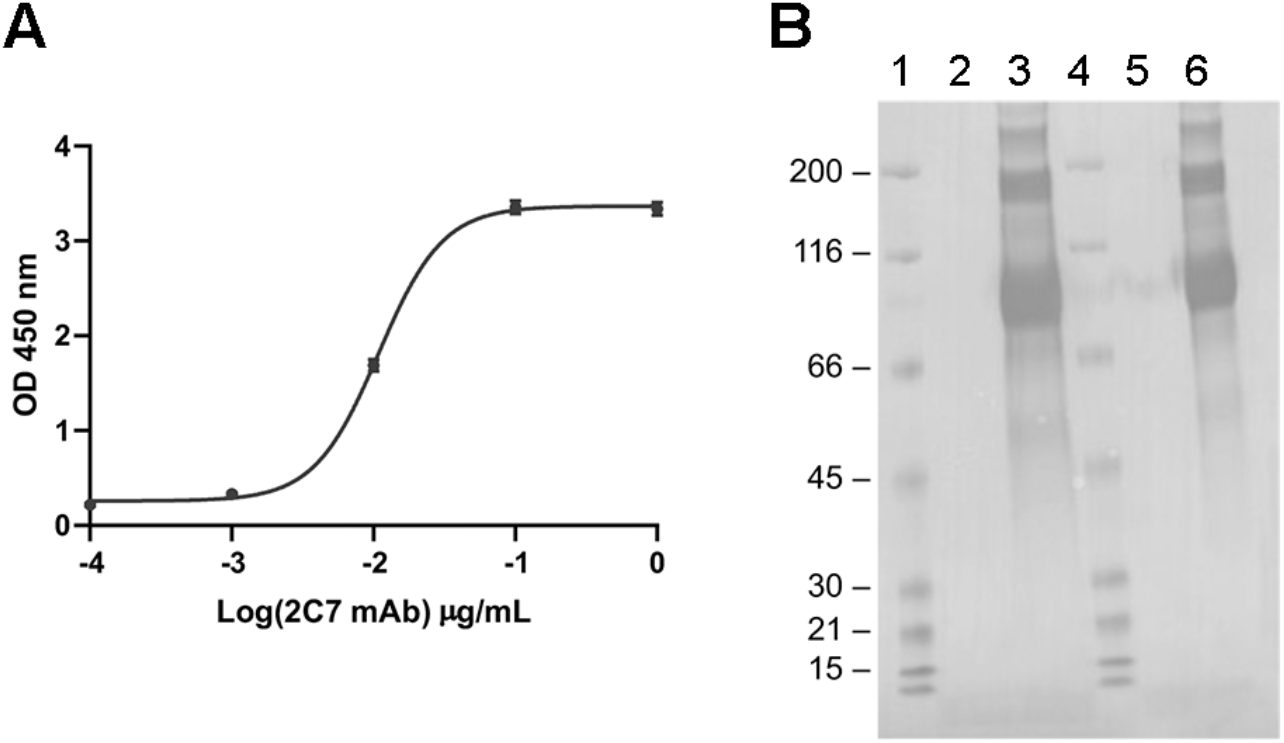
Binding of mAb 2C7 to tetrasaccharide-BSA conjugate. **A**, Concentration-dependent binding by ELISA. Each data point is the average of three measurements with the standard deviation plotted (SD for each data point was <10%). **B**, Western blot. Lane 1, Molecular weight (MW) markers (Bio-Rad); 2, BSA, 1.6 µg; 3, tetrasaccharide-BSA conjugate, 2 µg; 4, MW markers; 5, BSA; 6, tetrasaccharide-BSA conjugate, 1 µg. The masses of the MW markers in kDa are shown at the left. An average of 16 tetrasaccharides per BSA was conjugated based on MALDI-TOF MS analysis (**Fig. S3A**).

### Thermodynamics of octasaccharide and tetrasaccharide binding to mAb 2C7

We used isothermal titration calorimetry (ITC) to investigate the thermodynamic parameters of the mAb 2C7 interactions with the purified octasaccharide and synthetic tetrasaccharide. The data were fitted to a one-site binding model to derive thermodynamic parameters and dissociation constants. The thermodynamics for binding of the two glycans to mAb 2C7 were similar. The binding stoichiometries were 2.1 indicating that two molecules of each OS bound to one molecule of mAb (**Fig. 6**), which is expected for an IgG antibody with two antigen-binding sites [34]. The derived dissociation constants, *KD* (=1/*KA*), were 6.35 *μ*M for the octasaccharide and 1.99 µM for the tetrasaccharide, which indicated that the mAb has slightly higher affinity for the tetrasaccharide than the octasaccharide (**Fig. 6**). The interactions between mAb 2C7 and each OS were exothermic processes (Δ*G* ∼ -29.7 and -32.6 kJ/mol for octasaccharide and tetrasaccharide, respectively). Binding reactions had large, negative enthalpies (Δ*H* = −27.3 and -43.3 kJ/mol) and smaller entropic contributions (−*T*Δ*S* = −2.4 and +10.8 kJ/mol). Therefore, binding was enthalpically driven, which is consistent with the large number of H bonds in the complex. The observed entropic penalty for the tetrasaccharide suggests a loss of conformational freedom upon binding and/or the absence of stabilizing *π*-stacking interactions involving the α-chain in the synthetic tetrasaccharide.

**Figure 6.**
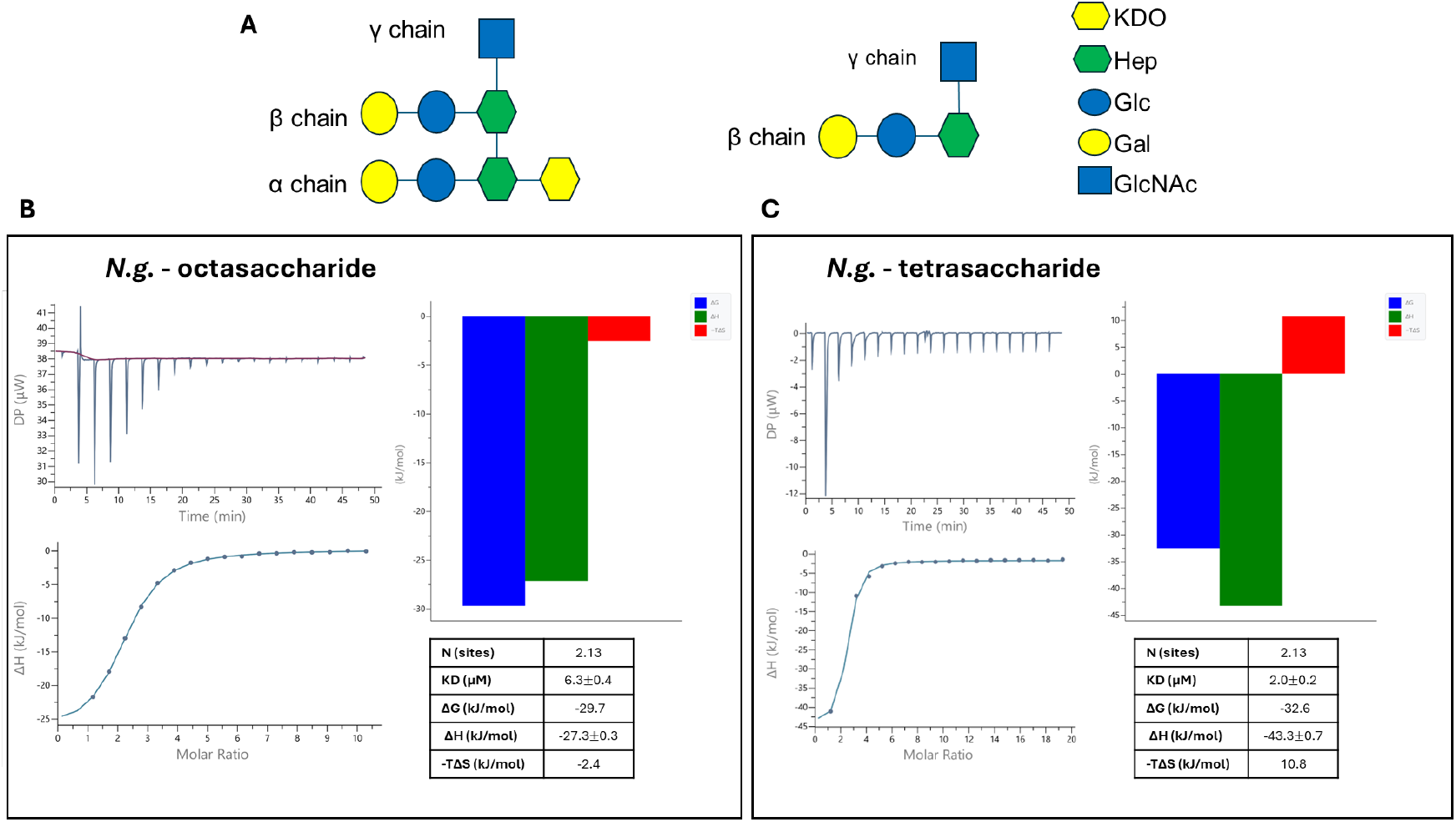
Thermodynamics of binding of purified octasaccharide and synthetic tetrasaccharide to mAb 2C7 by isothermal titration calorimetry. **A**, Schematic structures of octameric and tetrameric OS. **B**, ITC data and fit parameters for octasaccharide. Upper left, baseline-leveled raw data of mAb 2C7 titration with octasaccharide. Lower left, resulting curve fitted using a single binding site model. Upper right, histograms of the thermodynamic parameters defining the binding of octasaccharide to mAb 2C7. Lower right, summary of thermodynamic parameters. **C**, ITC data and fit parameters for tetrasaccharide. The layout of graphs is the same as in panel B.

### Bound conformation of tetrasaccharide and interactions with mAb 2C7

To assess whether key recognition features identified for the purified octasaccharide are retained in the synthetic tetrasaccharide, we investigated its interaction with mAb 2C7 using NMR spectroscopy and MD simulations. Ligand-based NMR approaches were employed to investigate the molecular details of the interaction, which allowed us to map the tetrasaccharide epitope, and therefore to identify the residues of the tetrasaccharide that comprised the binding interface with mAb 2C7 (**Fig. 7 A, B**). STD NMR revealed that mAb 2C7 primarily recognizes the β chain linked to HepII, particularly the terminal βGal residue. The strongest STD enhancements were observed for the H3 and H4 protons of the terminal βGal, with additional contributions from H2 and H5. Furthermore, the αGlc residue showed STD signals, particularly at H3, with a minor contribution from H2 and H4, suggesting a partial role in stabilizing the interaction. Within the γ-chain, the αGlcNAc exhibited the most prominent STD effect at H2, with lower contributions from H3 and H4. αHepII moiety displayed significant contributions at H2 and H3. Overall, these data indicate that the key recognition features identified for the octasaccharide (see section 3.2) are retained in the tetrasaccharide, with binding dominated by the β-γ-chains. To further investigate the bioactive conformation of the tetrasaccharide in the bound state (**Fig. 7D**), tr-ROESY experiments reveal no significant conformational changes between free and bound states.

**Figure 7.**
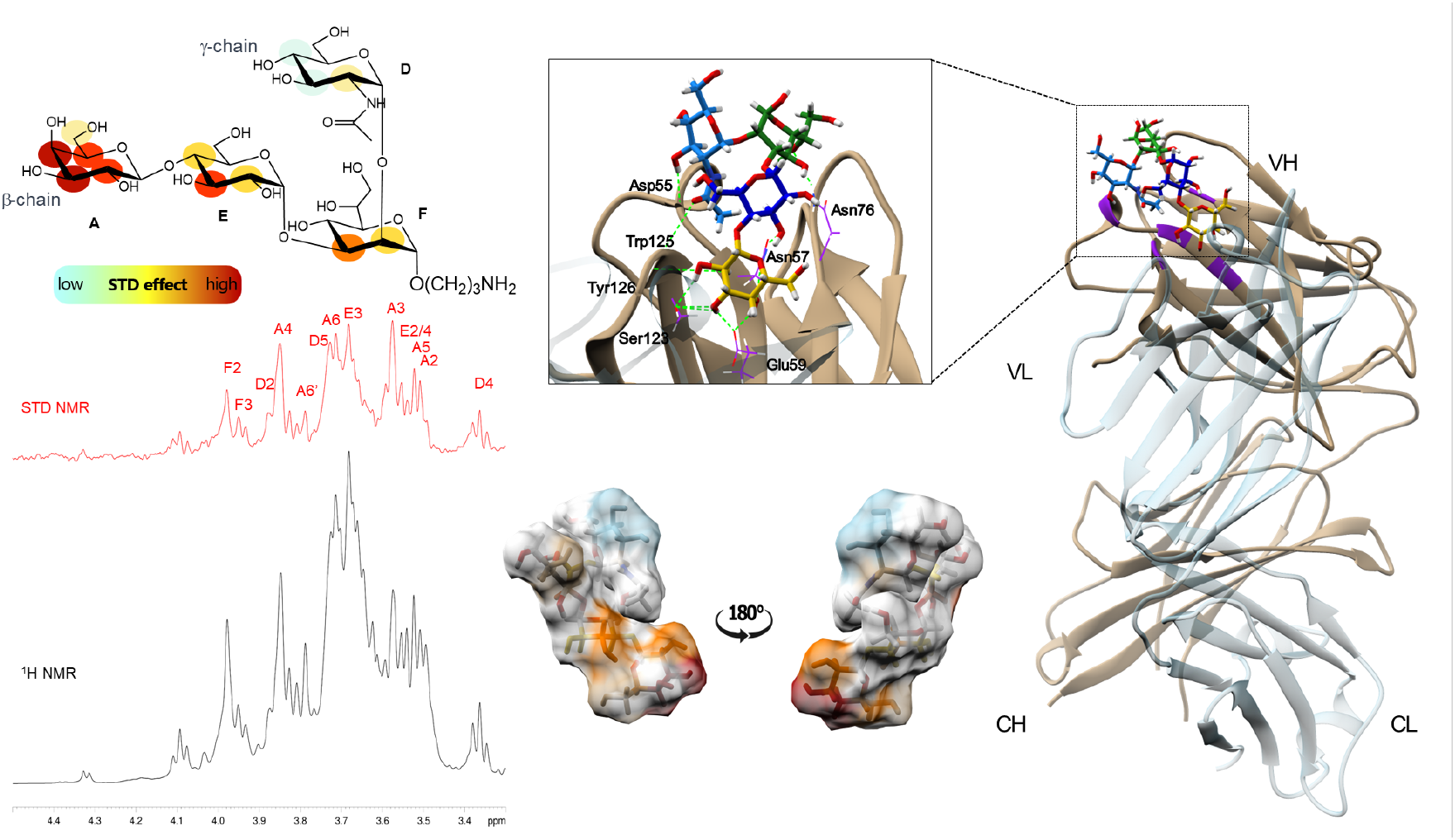
Epitope mapping of mAb 2C7-tetrasaccharide interactions. **A**, STD NMR analysis and epitope mapping of synthetic tetrasaccharide. Schematic view of tetrasaccharide structure colored according to STD values. **B**, ^1^H and STD spectra; %STD are obtained by the ratio (I_0_ − I_sat_)/I_0_, where (I_0_ − I_sat_) is the signal intensity in the STD-NMR spectrum (red) and I_0_ is the peak intensity of the off-resonance spectrum (black). **C**, Representative 3D model of the bioactive complex highlighting H-bond interactions involving the terminal βGal residue, which establishes recurrent contacts with the side chains of residues Glu59, Ser123, Tyr126, and Asn57. Fab CH and CL chains are shown in beige and light grey, respectively and the constituent domains variable heavy (VH), variable light (VL), constant heavy (CH) and constant light (CL) are labeled. **D**, Surface representation of the bioactive conformation of tetrasaccharide bound to Fab 2C7. The ligand is shown in its lowest-energy pose colored according to the STD intensity distribution.

As described above for the octasaccharide, we constructed a 3D model of the model of the mAb 2C7– tetrasaccharide complex and subjected it to a 500 ns MD simulation. We observed an excellent agreement between the experimental STD and computational MD experiments. Cluster analysis based on ligand RMSD showed good convergence, and a representative structure of the most populated cluster (**Fig. 7C**) confirmed a binding mode consistent with that observed for the purified octasaccharide.

Significant H-bond interactions were sampled, primarily involving the hydroxyl groups at positions 2, 3, and 4 of the terminal βGal. Specifically, βGal-O2 formed H bonds with Ser123 as an acceptor (17%) and with Tyr126 as a donor (47%). For terminal βGal-O3, interactions were observed with Glu59 and Ser123 as acceptors (54% and 16%, respectively), while Ser123 also acted as a donor (12%). At the βGal-O4 position, Glu59 was identified as a predominant acceptor (88%), and Asn57 contributed as a donor (44%).

## DISCUSSION

mAb 2C7 recognizes an epitope on gonococcal LOS that has many desirable attributes for a vaccine or therapeutic target. On a molar basis, it is the most abundant molecule on the surface of gonococci and is an important virulence factor. The 2C7 epitope is expressed by most clinical isolates, which suggests a role in pathogenesis or immune evasion [25]. mAb 2C7 elicits complement-dependent bactericidal activity against strains expressing the epitope [22, 25]. The mAb decreases colonization by gonococci in a mouse vaginal model of infection [16, 26]. Finally, unlike other LOS epitopes such as lacto-N-neotetraose (Gal-GlcNAc-Gal-Glc) and the P^*K*^-like epitope (Gal-Gal-Glc) [35], the 2C7 epitope does not mimic human glycans [25], which is an important consideration for a mAb 2C7 epitope-based vaccine or therapeutic.

Numerous previous studies have sought to define the epitope recognized by mAb 2C7 [24, 36-38]. These studies employed wild-type strains expressing different LOS structures [25], strains with phase variation in the *lgt* loci (e.g. MS11mk strains; [39]), or mutant strains with genetically stabilized LOS structures [24]. Given the importance of the 2C7 epitope in pathogenesis and as a potential vaccine or therapeutic target, in the present study we used chemical, biophysical and structural biology approaches to define the 2C7 epitope.

The crystal structure of the Fab 2C7-octasaccharide complex at 1.6 Å resolution shows numerous interactions between the Fab and glycan, which are identical in the two copies in the asymmetric unit of the crystal. The independent structure derived from NMR experiments and MD simulations highlights the dynamics of the complex and is in close agreement with the crystal structure. In both structures, the β and γ chains extending from HepII form a contiguous lobe that protrudes into a cleft of the Fab formed by the CDR loops. In the crystal structure, the β chain (lactose) contributes most of the buried surface area to the complex (53%), followed by the γ chain (24%) and the α chain (19%). In the MD structure, the α chain is dynamic and makes only transient contacts with the Fab (**Fig. S9**).

The structure of the Fab 2C7-octasaccharide complex explains the specificity of the mAb for different LOS α chain variants. mAb 2C7 binds strongly to LOS when the α chain includes two hexoses but less well to a variant with three, four or five hexoses [24]. In the crystal structure of the complex, the α chain lies on the surface of the Fab and interacts hydrophobically with an aromatic side chain. This conformation allows the Fab to bind LOS variants with longer α-chain extensions, however certain conformations of longer α chain extensions might interfere with mAb binding. In particular, the P^*K*^-like α-chain (Galα1-4Galβ1-4Glc) significantly reduced binding and bactericidal activity of mAb 2C7 [24, 38]. The α linkage of terminal βGal in the P^*K*^-like structure contrasts with the β linkages of all other hexoses of the α chain, which may spatially interfere with binding of mAb 2C7.

The crystal structure also shows a critical role of the β chain in the interaction with Fab 2C7, mediating multiple H bonds and substantial buried surface area. The mAb binds strongly to LOS when the β chain is lactose, but there is no binding when this chain is either absent or extended to four residues in length (13,40). An *lgtE* deletion mutant of strain 15253, which elaborates only a Glc residue from both HepI and HepII, does not bind mAb 2C7 [37]. These observations underscore the importance of the terminal βGal of the β chain for the mAb 2C7-LOS interaction. Previous studies of gonococci that expressed lactose from HepI and HepII grown in media containing CMP-Neu5Ac (25 µg/mL), which results in sialylation of the β chain of LOS, decreased binding of the mAb by 22-55% [22]. In the structure of the Fab 2C7-OS complex, there is insufficient space in the cleft of the Fab to accommodate a sialic acid residue on the β chain. We previously showed that growth of gonococci in the presence of CMP-Neu5Ac (20 µg/mL) resulted in partial LOS sialylation [40]. Therefore, it is likely that gonococci grown in media containing a similar concentration of CMP-Neu5Ac contained both sialylated and non-sialylated species, and that mAb 2C7 bound to the non-sialylated LOS molecules [22].

Based on the structure, the γ chain (GlcNAc) plays a key role in the epitope recognized by mAb 2C7. This chain is added to HepII by the enzyme RfaK prior to the addition of the proximal Glc residues of the α and β chains [41]. Deleting *rfaK* results in expression of LOS devoid of α and β chains. Although not experimentally tested with *N. gonorrhoeae*, an *rfaK* deletion mutant would not be predicted to bind to mAb 2C7. Thus, prior to the current study, it was not possible to assess the importance of the γ chain in mediating binding to mAb 2C7 by genetically manipulating LOS. It is noteworthy that harsh alkaline hydrolysis (4 M KOH, 4 h, 120 °C) of 15253 LOS to completely de-acylate LOS abrogated mAb 2C7 binding [37]. This treatment is expected to also de-N-acetylate GlcNAc, thereby providing evidence for the importance of the acetyl moiety of the γ chain GlcNAc in binding mAb 2C7.

In previous studies, we identified a peptide that mimics the 2C7 epitope and determined the crystal structure of the peptide bound to Fab 2C7 [28]. A superposition of Fab 2C7 in the peptide and octasaccharide complexes show that the two ligands are bound in the same cleft and primarily interact with the heavy chain (**Fig. S11**). Further, several of the same heavy chain residues, N57, N76 and W125, form H bonds with both ligands (**Table S7**). In the peptide complex, Phe12 is the most buried residue, whereas in the octasaccharide complex, the βGal residue in the β chain is most buried. Both ligand residues interact hydrophobically with heavy chain residue Y126. Some key differences between the binding modes of the two ligands are that the peptide interacts with the Fab predominantly through hydrophobic and aromatic interactions, whereas the octasaccharide interacts with the Fab via direct H bonds involving β- and γ-chain residues. Thus, while the binding pockets of the Fab are similar in the two complexes, the chemical nature of the interactions is distinct.

The crystal structure of Fab 2C7 bound to purified octasaccharide provides the first high-resolution structural information on gonococcal LOS. A crystal structure of a meningococcal LOS bound to Fab LPT3-1 at 2.7-Å resolution was reported previously [42]. While the two LOS variants share the same core structure (KDO, HepI and HepII) and the same γ chain (GlcNAc), in the meningococcal OS the α chain was Glc and the β chain was absent. Superposition of the Fab variable regions shows that OS from the two species bind with the sugars at the non-reducing ends (α, β and/or γ chains) oriented toward the Fab, and the core KDO and Hep residues are oriented toward the solvent (**Fig. S12**). In the meningococcal LOS complex, the binding specificity is primarily driven by the γ chain, while in the gonococcal octasaccharide complex, the specificity derives from both the β and γ chains, which form a contiguous lobe. In both complexes, aromatic residues within the antibody binding site form stacking or hydrophobic interactions with the carbohydrate residues. However, whereas the meningococcal antibody recognizes a trisaccharide epitope, Fab 2C7 binds a distinct tetrasaccharide epitope.

Finally, our structural and binding data can guide the future development of vaccines and therapeutics based on the conserved 2C7 epitope. The structure of the Fab 2C7-octasaccharide complex can be used to humanize mAb 2C7 variable regions to optimize an immunotherapeutic with minimal anti-mouse antibody responses. Specifically, the structure identifies both the paratope residues in contact with the octasaccharide ligand and second shell (Vernier zone) residues that are likely important for the conformation of the paratope. Complementary immunological and biophysical approaches showed that a tetrasaccharide comprising HepII and its β and γ extensions represents a minimal 2C7 epitope, which could form the basis of a vaccine. Further studies are warranted to conjugate tetrasaccharide or octasaccharide glycans to a suitable carrier protein to develop a broadly protective gonococcal vaccine.

## EXPERIMENTAL

### Materials

For NMR studies, we used phosphate-buffered saline tablets (Sigma) dissolved in deuterated water. For crystallization experiments, we employed sparse matrix screening reagents (Hampton Research) and molecular biology grade chemicals (Sigma, Fluka and Hampton Research). No unexpected or unusually high safety hazards were encountered.

### Extraction and purification of LOS from gonococcal strain 15253

Dried bacterial cells (5 g) were extracted using the hot phenol/water method [29]. Sodium dodecyl sulfate-polyacrylamide gel electrophoresis (SDS-PAGE; 13.5%) with silver staining [30] showed the presence of LOS in the aqueous phase. The aqueous phase was treated with RNase and DNase (Roth) at 37 °C overnight, and proteinase K (Roth) at 57 °C for 4 h, followed by dialysis (molecular weight cut-off 12-14 kDa) against distilled water for 48 h. For SDS-PAGE, the purified LOS was prepared at a concentration of 1 mg/mL, boiled for 10 minutes, and 4 to 8 µg were loaded into wells of a 13.5% SDS polyacrylamide gel with a 5% stacking gel. After removal of nucleic acids and proteins, dialysis, centrifugation, and purification by gel filtration chromatography), the LOS was washed with chloroform–methanol (1:2 v/v) and chloroform–methanol–water (3:2:0.25 v/v) mixtures to remove residual phospholipids. Purity of the preparation was again assessed by SDS–PAGE.

### Monosaccharide analysis of *Neisseria gonorrhoeae* LOS

The monosaccharide composition of *N. gonorrhoeae* LOS was determined by analyzing the acetylated O-methylglycoside derivatives resulting from treatment with 1.25 M HCl in methanol at 100 °C, followed by acetylation with acetic anhydride in pyridine at 85 °C for 30 minutes. All chemical analyses were carried out using gas-liquid chromatography with an Agilent Technologies 6850A system equipped with a 5973N mass-selective detector and a Zebron ZB-5 capillary column (Phenomenex, 30 m × 0.25 mm inner diameter, 0.25 μm film thickness). Helium was used as the carrier gas at a flow rate of 1 mL/min. The program for sugar analysis was as follows: an initial temperature of 150 °C held for 5 minutes, followed by a ramp from 150 °C to 280 °C, at 3 °C/min, and a final hold at 280 °C for 5 minutes.

### Separation of OS from lipid A

An aliquot (100 mg) of LOS sample was subjected to mild acetic acid hydrolysis with 1% acetic acid at 100 °C for 5 hours to cleave the bond between the lipid A and OS portions. After hydrolysis, the mixture was centrifuged (7000 × *g*, 60 min, 4 °C) to separate the insoluble lipid A fraction from the soluble OS. The supernatant containing the OS was collected separately and lyophilized. The second Kdo is removed by the mild acid treatment. The dried material was purified by gel filtration chromatography using Bio-Gel TSK 40 to remove contaminating lipid A and reaction products.

### MALDI-ToF MS analysis

Linear and Reflectron MALDI-TOF MS spectra of LOS mild hydrolysis product (OS) were analzyed with an Applied Biosystems AB SCIEX TOF/TOF 5800 mass spectrometer equipped with an Nd:YAG laser (λ = 349 nm), with a 3-ns pulse width and a repetition rate of up to 1000 Hz. The OS sample was prepared by dissolving an aliquot in distilled water and mixing it with 2,4,6-trihydroxyacetophenone (THAP) in a solution of acetonitrile with 0.1%TFA (1:1). Each spectrum in the MS experiments was a result of the accumulation of 2000 laser shots. Each experiment was performed in duplicate.

### NMR spectroscopy

#### Instrument and sample conditions

All NMR experiments were recorded at 298 K on a Bruker AVANCE NEO 600-MHz spectrometer equipped with a cryo-probe. Data acquisition and processing were performed using TOPSPIN 4.1.1 software. Samples were prepared in phosphate-buffered saline (10 mM Na2HPO4, 2.7 mM KCl, 137 mM NaCl, 10 mM NaN3, pH 7.4) at 298 K, using 0.05 mM sodium 3-(trimethylsilyl) propionate-d4 (TSP) as the internal reference.

#### Structure determination experiments

Total correlation spectroscopy (TOCSY) experiments were performed with spinlock times of 100 ms and data set (t1×t2) of 4096×800 points. The experiments of Rotating-frame Overhauser enhancement spectroscopy (ROESY) and nuclear Overhauser enhancement spectroscopy (NOESY) were performed with data sets (t1×t2) of 4096×800 points and mixing times ranging from 100 to 400 ms. Double-quantum-filtered phase-sensitive correlation spectroscopy (DQF-COSY) experiments were recorded using data sets of 4096×912 points. Heteronuclear single-quantum coherence (HSQC) and heteronuclear multiple-bond correlation (HMBC) experiments were carried out in the ^1^H-detection mode using single-quantum coherence with proton decoupling in the ^13^C domain using data sets of 2048×512 points. Phase-sensitive mode and sensitivity improvements were applied for HSQC, along with echo/antiecho gradient selection and multiplicity editing in the selection step. In the HMBC experiments, an optimization was applied for long-range coupling constants, incorporating a low-pass J filter to eliminate the one-bond correlations, via gradient pulses for selection.

#### Ligand-based NMR experiments

Saturation transfer difference (STD) NMR experiments were conducted with a 1:50 protein-ligand ratio. STD NMR spectra were acquired using 32k data points, zero-filled to 64k before processing. Protein resonances were selectively irradiated with 40 Gauss pulses lasting 50 ms, with the off-resonance pulse frequency set at 40 ppm and the on-resonance pulses at 0 ppm. To suppress protein signals, a 20 ms spinlock pulse was applied. The STD NMR experiments were performed at a saturation time of 2 seconds. Ligand epitope mapping was achieved by calculating the ratio (I0 – Isat)/I0, where Isat corresponds to the intensity of the STD NMR signal and I0 to the intensity of the off-resonance spectrum. The strongest STD response was normalized to 100%, and all other proton STD signals were scaled accordingly. Control experiments were also performed in the absence of the protein to ensure the specificity of the observed STD signals. Transferred NOESY (tr-NOESY) experiments were also conducted with a 1:50 protein-ligand ratio. Spectra were recorded with data sets of 800 points, using a mixing time of 100-300 ms.

### Crystallization and X-ray structure determination

Chimeric Fab 2C7 was expressed in CHO cells and purified by Protein G affinity chromatography by a commercial vendor (Genscript). The Fab was concentrated to 6.0 mg/ml and mixed with a 10-fold excess of octasaccharide, prepared as described above (see section 2.4). Crystals of the complex were obtained in Crystal Screen Cryo solution #40 (Hampton Research) without further optimization. The solution contains 0.095 M sodium citrate tribasic dihydrate, pH 5.6, 19% v/v 2-propanol, 19% w/v polyethylene glycol 4,000 and 5% glycerol. The crystals were mounted on nylon loops, flash cooled in liquid N2 and X-ray data were collected on beam line 8.3.1 at the Advanced Light Source, Lawrence Berkeley National Laboratory. We collected 360° of oscillation data with a Δϕ of 0.5°, an exposure time of 0.05 sec, a detector distance of 250 mm and a wavelength of 1.116 Å. The data collection and refinement statistics are listed in **Table S1**. The structure of the complex was solved by molecular replacement using the structure of chimeric Fab 2C7 alone (PDB ID 8DOZ) [28] and Phaser-MR in the Phenix software package [43]. The crystals were in space group C2 and there were two copies of the complex in the asymmetric unit of the crystal. After initial refinement of the MR solution at 1.6-Å resolution, there were prominent electron density features consistent with bound octasaccharide. We built the model containing two copies each of Fab and octasaccharide, as well as solvent in Coot [44]. The model was refined with phenix.refine using torsional non-crystallographic symmetry restraints throughout the refinement. Iterative rounds of model building and refinement resulted in a model with good geometry, an R-factor of 0.189 and a free R-factor of 0.209. The structure was analyzed using PyMol (The PyMOL Molecular Graphics System, Version 3.0 Schrödinger, LLC.) and the Proteins, Interfaces, Structures and Assemblies (PISA) server [45]. The atomic coordinates and structure factors have been deposited in the Protein Data Bank (accession code 9Y5C).

### Computational studies

Standard monosaccharide structures were downloaded from the GLYCAM website (http://www.glycam.org) [46]. Non-standard residues, including KDO, HepI, GlcN and HepII, were parametrized using a custom protocol involving the calculation of Restrained Electrostatic Potential (RESP) charges from Hartree-Fock calculations with a 6–31G* basis set. The Antechamber and xLeap modules were used to generate .prep and .frcmod files [47]. The parameters generated in this work have since been incorporated into the GLYCAM database and are currently available through the GLYCAM website [48].

Molecular mechanics (MM) calculations were carried out using the MM3* force field in a vacuum environment, with a dielectric constant of 80. For the disaccharide structures, the glycosidic torsion angles (φ and ψ) were systematically varied in 18° increments. At each (φ, ψ) point on the map, optimization was performed using 2000 P.R.

Molecular dynamics (MD) simulations were performed by using AMBER18 software package [49], employing explicit water molecules. The AMBER ff14SB, Glycam06j-1, and TIP3P force fields were used for the protein residues, OS ligand, and solvent, respectively. Preparation of Fab 2C7 [28] involved adding missing hydrogen (H) atoms, determining the protonation states of ionizable groups, and capping termini using the Maestro Protein Preparation Wizard [50]. Each system was hydrated with an octahedral TIP3P water box, buffered to 15 Å, and neutralized with counterions. Input files were generated using the *tleap* module in AMBER. Minimization steps were carried out using the Sander module, while MD simulations were performed with the *pmemd* module. The initial structure was refined through energy minimization, applying the SHAKE algorithm to constrain C-H bonds and using a 1 fs integration step. Periodic boundary conditions were applied, and electrostatic interactions were modeled with the smooth particle mesh Ewald method, using a grid spacing of 1 Å. Minimization was performed in two stages: first, the complex was restrained while the solvent was minimized, followed by minimization of the entire system. The system was then gradually heated from 0 K to 300 K. Heating was conducted in two steps: from 0 K to 100 K under constant volume and from 100 K to 300 K under isobaric conditions. Once the target temperature of 300 K was reached, the system was equilibrated over 50 ps with progressive energy minimization and solute restraints. After equilibration, restraints were removed, and production runs were performed under isothermal-isobaric conditions. Coordinates were saved throughout the 500 ns production simulations, with trajectory snapshots recorded every 2 ps. Trajectories were analyzed with the ptraj module in AMBER 18, and the MD results were visualized using the VMD program [51]. Cluster analysis of the trajectories was performed based on the ligand RMSD, using the K-means algorithm implemented in *ptraj*. The representative structure of the most populated cluster was selected to characterize the interactions within the complexes. H bonds were identified using the CPPTAJ module in AMBER18 [47], with a H bond defined by a donor-acceptor distance cutoff of 3 Å and an A-H-D angle cutoff of 135°. Three-dimensional images of the complexes were generated using UCSF Chimera [52] and PyMol (The PyMOL Molecular Graphics System, Version 3.0 Schrödinger, LLC.). Dihedral angle conformations were analyzed with a custom script, which depicted torsional variations during the MD simulations and generated histograms showing the most populated torsion values.

### Chemical synthesis of tetrasaccharide and conjugation to bovine serum albumin (BSA)

We produced a tetrasaccharide comprising HepII and the β and γ chains via chemical synthesis incorporating an N-hydroxysuccinimide (NHS) linker (**Fig. S2**). The tetrasaccharide was conjugated to BSA via the linker by incubation in potassium phosphate buffer (pH 7.0). The average number of tetrasaccharides per BSA was determined from MALDI-TOF MS analysis (**Fig. S3A**). The tetrasaccharide-BSA conjugate was analyzed for size and purity by SDS-PAGE (4-12% polyacrylamide in MOPS buffer; Thermo Scientific) followed by Coomassie blue staining (**Fig. S3B**).

### Binding of mAb 2C7 to synthetic tetrasaccharide-BSA conjugate

For ELISA, a 96-well microtiter plate was coated with tetrasaccharide-BSA conjugate (10 μg/mL) in carbonate buffer (0.05 M NaHCO_3_/Na_2_CO_3_, pH 9.6) and incubated at 4 °C overnight. The plates were washed four times with PBST (PBS containing 0.5% Tween-20) and then blocked with 1% BSA in PBS at room temperature for 1 h. After washing with PBST, serial dilutions of mAb 2C7 in 0.1% BSA/PBS were added to each well. The plates were incubated at 37 °C for 2 h and washed four times with PBST. Horseradish peroxidase (HRP)-conjugated anti-mouse IgG secondary antibodies (Jackson ImmunoResearch) diluted 1:2000 in 0.1% BSA/PBS were then added to the wells and incubated at 37 °C for 1 h. After washing with PBST, the plates were developed by adding 3,3′,5,5′-tetramethylbenzidine (TMB) substrate. After incubation for 15 min at room temperature, the reaction was quenched with 0.5 M H_2_SO_4_. The optical density (OD) was measured at 450 nm using a microplate reader. Data were analyzed using GraphPad Prism and values are presented as means ± SD (n=3).

For Western blotting, the BSA-tetrasaccharide conjugate and BSA alone in LDS sample buffer (Thermo Scientific) containing 2-mercaptoethanol were run on a 4-12% Bis-Tris gel with MOPS running buffer (Thermo Scientific) and proteins were transferred to a PVDF membrane. The membrane was blocked with PBS-1% dry-milk for 1 h and then incubated with cell culture supernatant containing murine mAb 2C7 for 15 h at 4 °C. Bound mAb 2C7 was detected with anti-mouse IgG conjugated to alkaline phosphatase (Sigma; 1:1000 in PBS-0.05% Tween 20) and developed using 5-bromo-4-chloro-3-indolyl phosphate / nitro blue tetrazolium (BCIP/NBT) substrate.

### Isothermal titration calorimetry

Isothermal titration calorimetry (ITC) experiments were carried out at 25 °C by using a MicroCal PEAQ-ITC instrument (Malvern). mAb and octasaccharide were both dialyzed against PBS pH 7.4 to avoid buffer mismatch between solutions. A solution of mAb (30μM) was loaded into the cell, and for each measurement, octasaccharide or tetrasaccharide was added through a micro syringe to the antibody solution stirred at a speed of 750 rpm. The titration was performed by adding 2 μL from a 1.6 mM of octasaccharide or 0.8 mM of tetrasaccharide for 19 injections using a time duration of 150s. The data were analyzed and fitted to a one-binding site model, in which standard thermodynamic parameters of equilibrium were calculated by the following equation [53]:

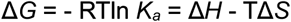

where Δ*G* is Gibbs free energy change, *K*_*a*_ is the equilibrium association constant, R is the gas constant, T is the temperature in kelvin, Δ*H* is the enthalpy change, and Δ*S* is the entropy change.

## SUPPORTING INFORMATION

Supplemental Tables: 7

Supplemental Figures: 12

## ACKNOWLEDGEMENTS

This work was supported by research grants from the National Institute for Allergy and Infectious Diseases, National Institutes of Health, R43 AI186979 (to S.R.), and R01AI182419 and R01AI190348 to X.H. X-ray data were collected at beamline 8.3.1 at the Advanced Light Source (ALS), Lawrence Berkeley National Laboratory. ALS is supported by the Director of the Office of Science, Office of Basic Energy Sciences, of the U.S. Department of Energy under Contract No. DE-AC02-05CH11231.

