## Supporting Information for "Structural basis for recognition of a gonococcal lipooligosaccharide epitope by monoclonal antibody 2C7 informs vaccine and immunotherapeutic design"

*BioRxiv* (submitted May 28, 2026)

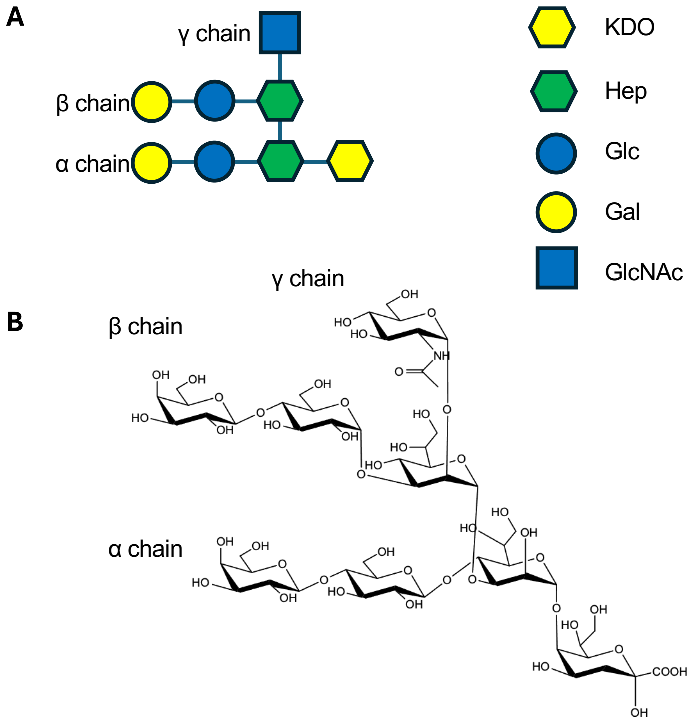

**Figure S1.** *Neisseria gonorrhoeae* strain 15253 oligosaccharide structure. **A,** Schematic representation using Symbol Nomenclature for Glycans. The legend is shown at the right. **B,** Haworth projection with reducing end KDO at bottom right. In both panels, the glycan extensions are labelled.

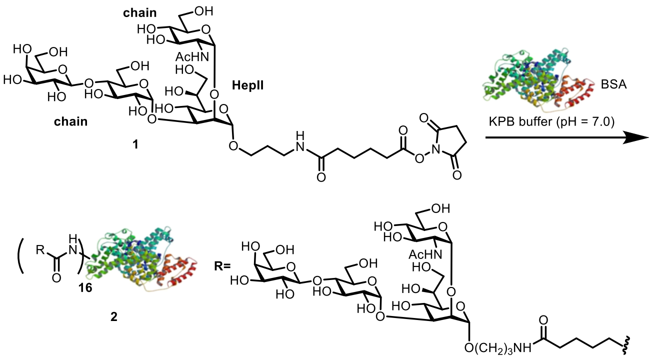

**Figure S2.** Synthesis of tetrasaccharide-BSA conjugate bearing the putative 2C7 epitope.

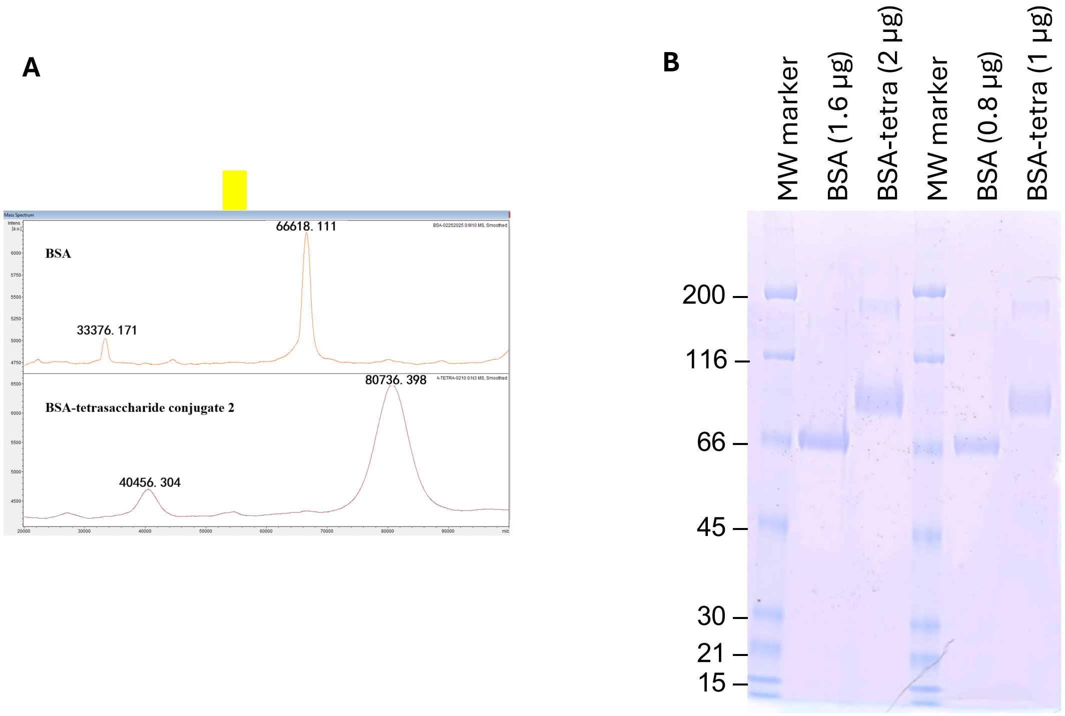

**Figure S3.** Characterization of tetrasaccharide-BSA conjugate. **A,** MALDI-TOF mass spectrometry. Based on the average molecular weight difference between the conjugate and native BSA, the average number of tetrasaccharides per BSA was calculated to be 16. **B,** SDS-PAGE. The tetrasaccharide conjugate and native BSA control were separated on a 4-12% NuPAGE gel in MOPS buffer (Thermo Scientific) and stained with Coomassie brilliant blue.

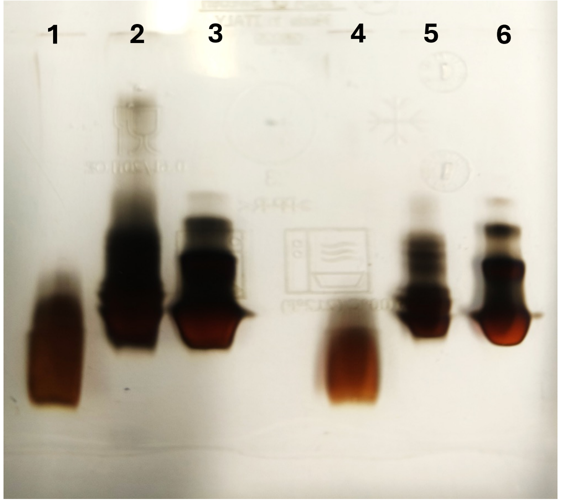

**Figure S4.** SDS-PAGE of gonococcal LOS. LOS was extracted with the hot phenol/water protocol and purified by SEC. LOS was separated by SDS-PAGE (13.5%) followed by staining with silver nitrate. Samples were:

1. *E. coli* LPS L-type (4 µg)
2. *N. gonorrhoeae* strain FA1090 wild-type LOS (8 µg)
3. *N. gonorrhoeae* strain 15253 LOS (8 µg)
4. *E. coli* LPS L-type (2 µg)
5. *N. gonorrhoeae* strain FA1090 (4 µg)
6. *N. gonorrhoeae* strain 15253 (4 µg)

| | | |
|---|---|---|
| **OSA** | **KDO; HepI; HepII; PEtN; Glc; Gal; Glc; Gal; GlcNAc** | **1595.5** *m/z* |
| **OSC** | **KDO; HepI; HepII; PEtN; Glc; Gal; Glc; Gal; GlcNAc; OAc** | **1637.5** *m/z* |

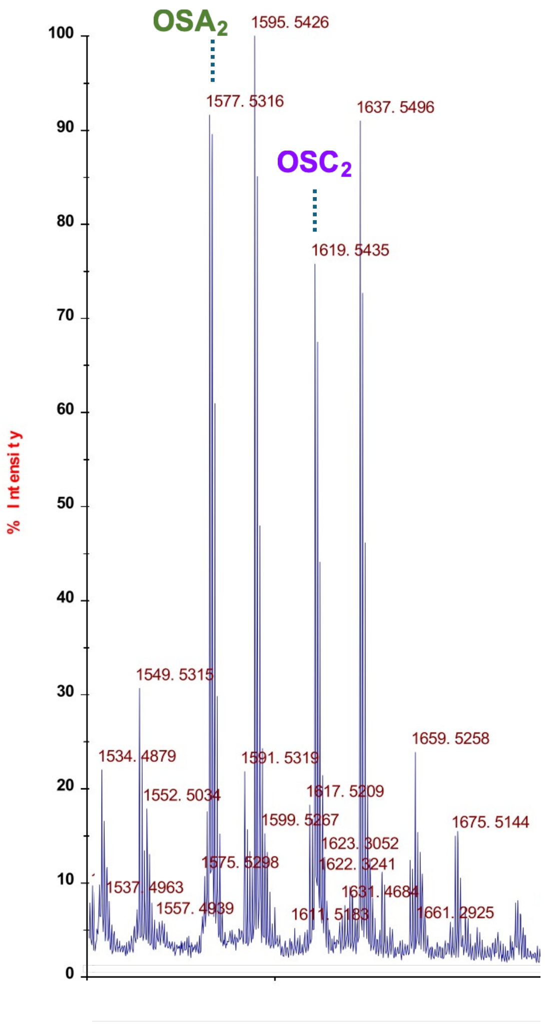

**Figure S5.** MALDI-TOF mass spectrum of the OS fraction obtained after mild acid hydrolysis of the LOS. Structural annotations are based on high-resolution accurate mass data and supported by compositional analysis. The OSA at *m/z* 1577.5316 contains 1 KDO, 2 Hep (heptose), 4 Hex, 1 HexNAc, and 1 PEtN (phosphoethanolamine). The ion at *m/z* 1637.5496 was attributed to OSC, composed of 1 KDO, 2 Hep, 4 Hex, 1 HexNAc, 1 PEtN, and 1 Ac (acetyl group).

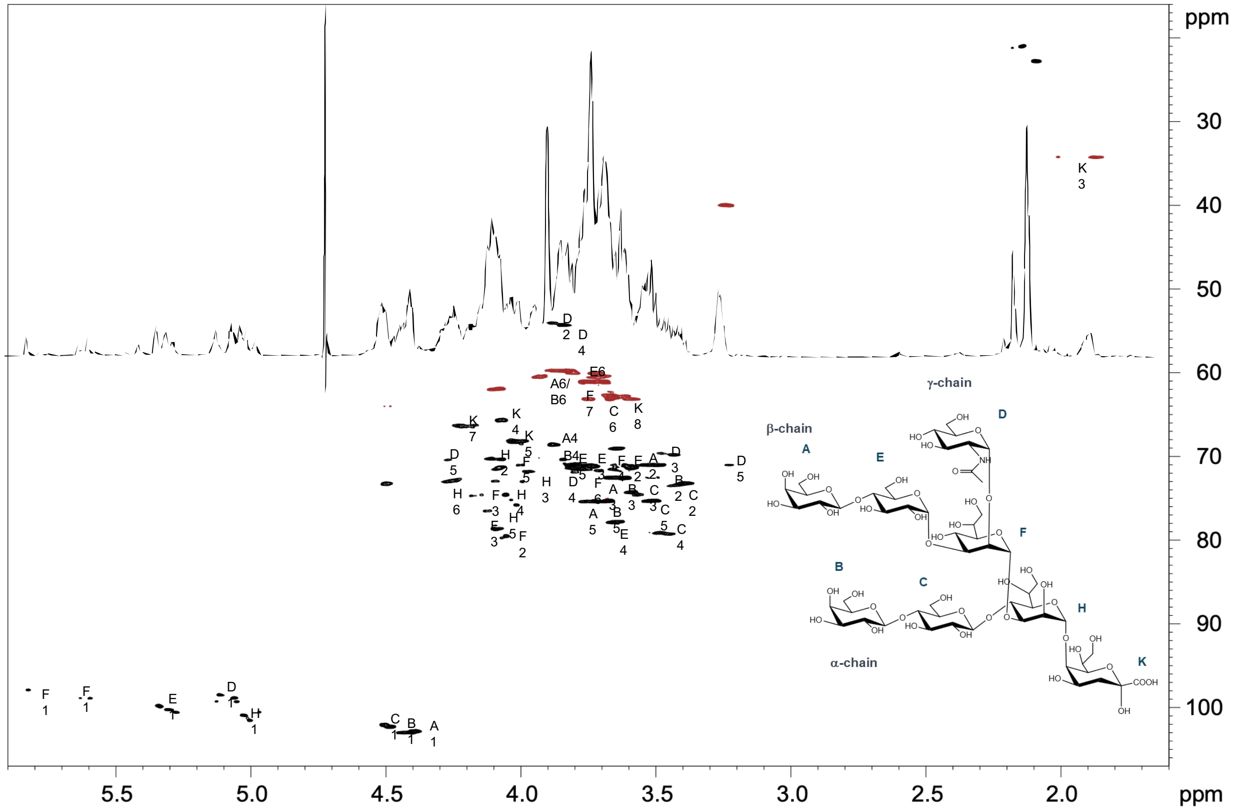

**Figure S6.** $^{1}H$-$^{13}C$- HSQC spectrum of purified octasaccharide from *N. gonorrhoeae* strain 15253.

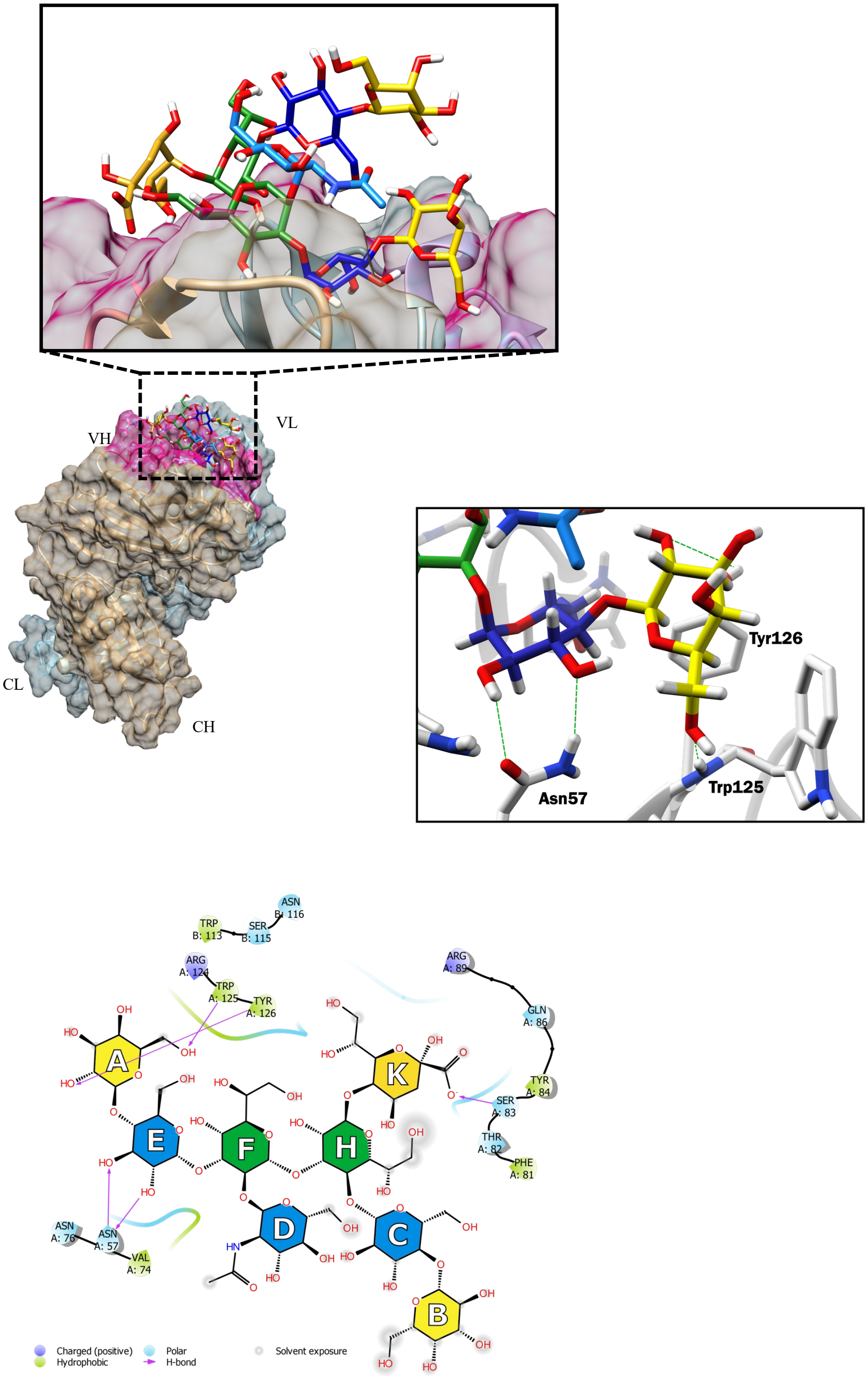

**Figure S7.** 3D structure of the Fab 2C7-octasaccharide complex that was used for MD simulations.

A

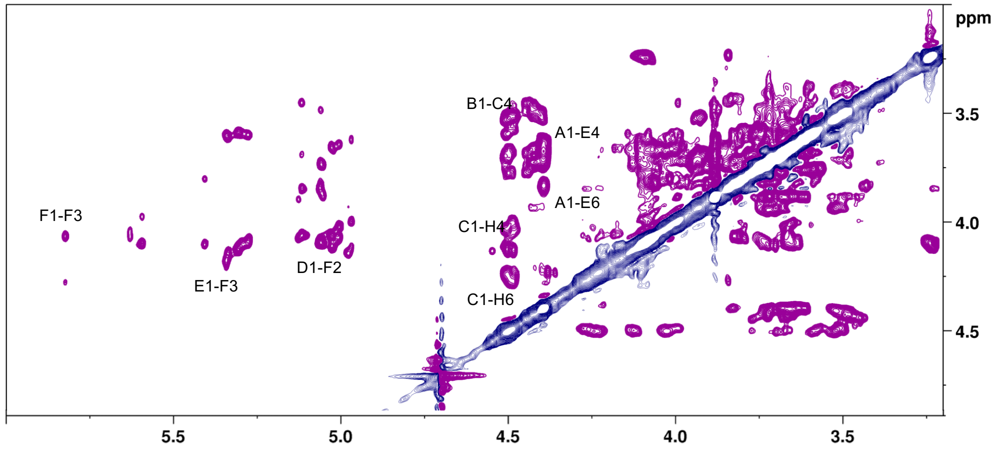

B

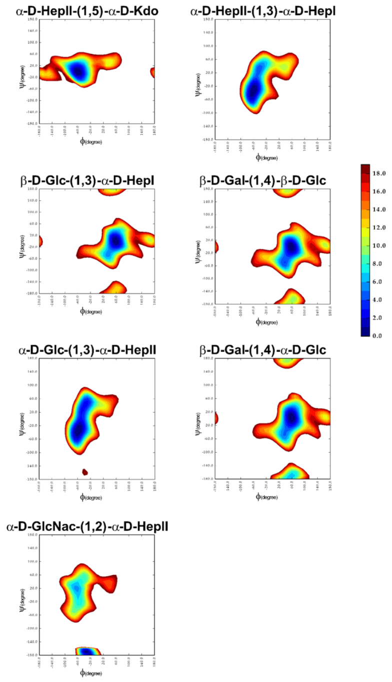

**Figure S8. A,** Transferred-ROESY NMR spectrum confirming intermolecular NOE contacts between OS protons and mAb2C7. **B,** Resulting adiabatic energy maps, which depict the φ and ψ glycosidic torsion angles

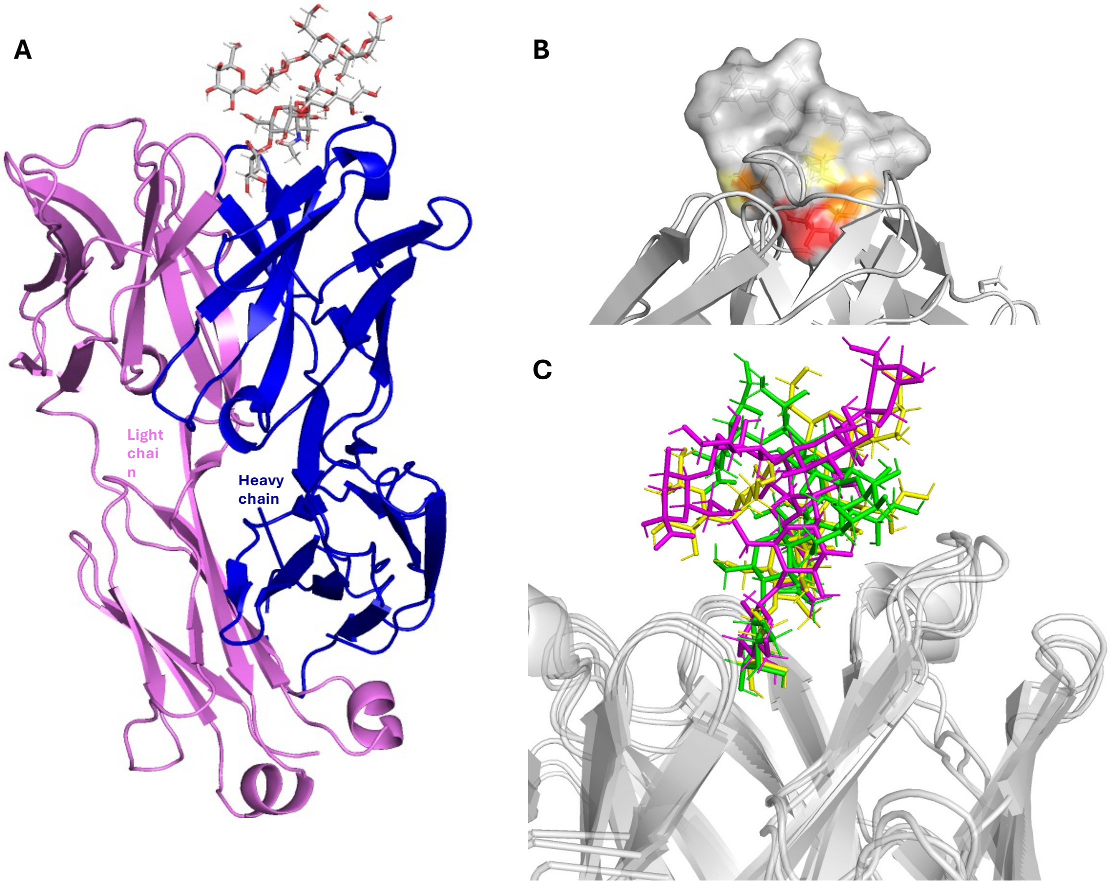

**Figure S9.** 3D views of Fab 2C7-octasaccharide complex obtained by NMR and MD. **A,** The most populated conformation obtained from the MD cluster analysis. **B,** Octasaccharide in its bioactive conformation, with the surface colored according to the STD effects (from the highest in red to the lowest in yellow), in complex with Fab 2C7. **C,** Superimposition of the different poses of Fab 2C7 in complex with octasaccharide in its bioactive conformation obtained from MD simulation.

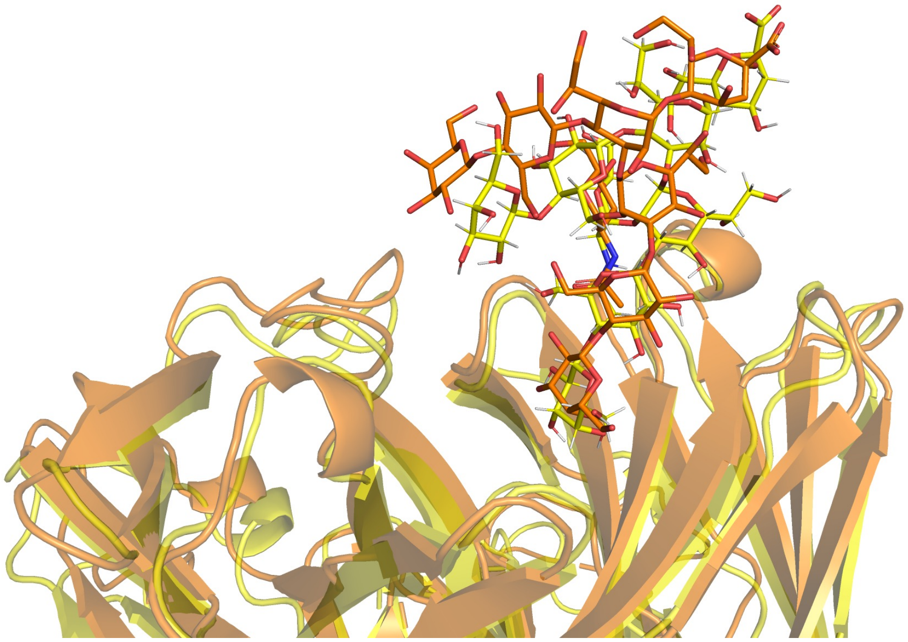

**Figure S10.** Representative MD conformation of Fab 2C7 in complex with the octasaccharide in its bioactive conformation as determined by NMR analysis (orange) superimposed onto the X-ray structure (yellow).

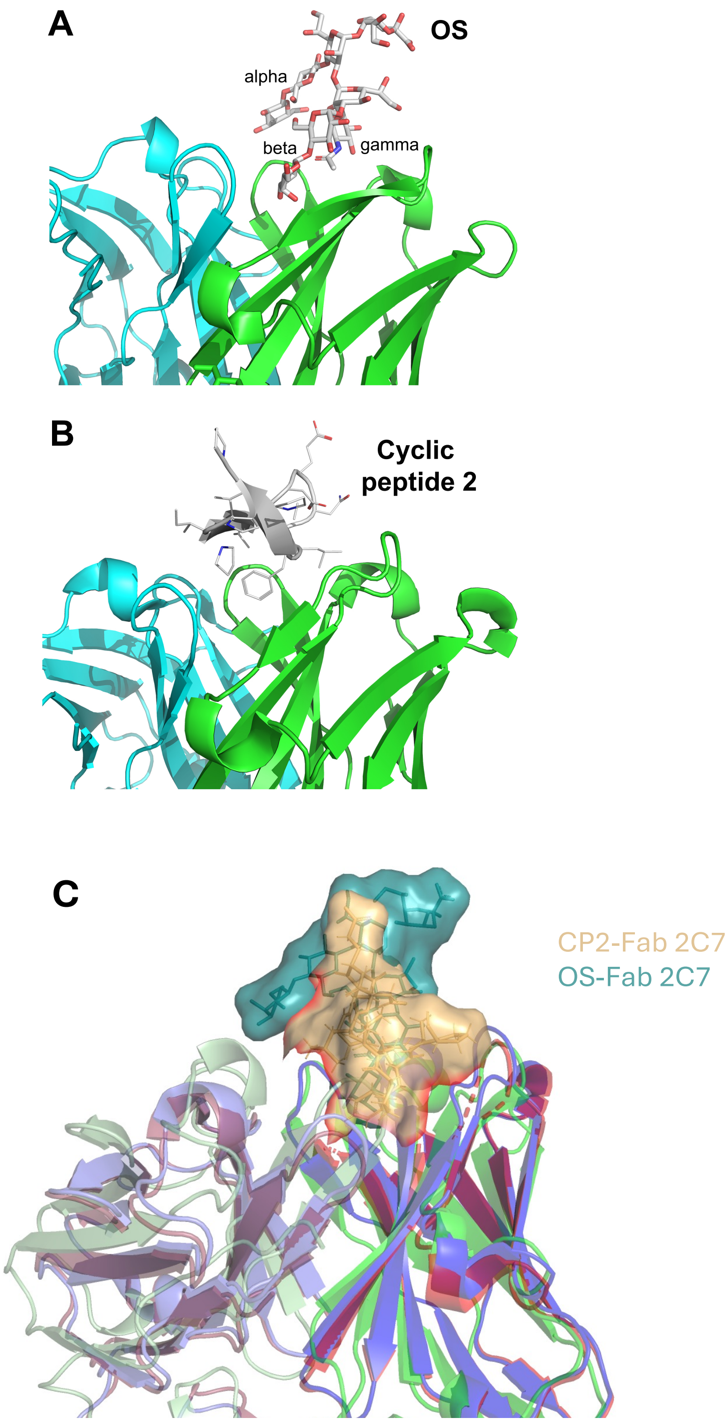

**Figure S11. A, B.** Comparison of binding modes of *N.gonorrhoeae* octasaccharide (**A**) and cyclic peptide 2 (CP2) (**B**), each bound to chimeric Fab 2C7. Coordinates for the latter structure were from PBD ID 8DUZ (Beernink et al. JACS Au 2024). The structures were superimposed based on Fab backbone atoms using PyMol 3.1 (https://pymol.org). Heavy chains are shown in green and light chains are shown in aqua. **C,** Superposition of the apo- form of Fab 2C7 (PDB ID 8DOZ) (purple color), Fab 2C7 complex with mimetic peptide (8DUZ) (red color), and Fab 2C7-OS complex (PDB ID 9Y5C) (green color). The mimetic peptide is shown with orange color and the OS with blue color. The coordinates for 8DOZ and 8DUZ were reported previously (Beernink et al. JACS Au 2024).

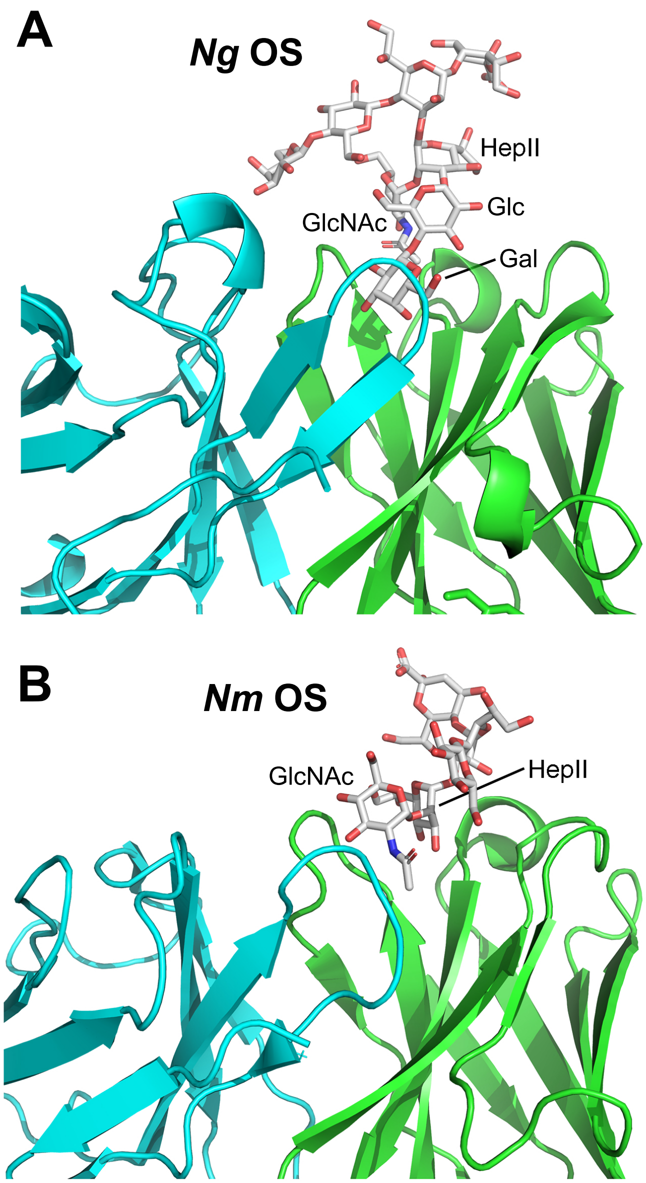

**Figure S12.** Comparison of binding modes of gonococcal OS to Fab 2C7 (**A**) and meningococcal OS to Fab LPT3-1 (**B**). Coordinates for latter structure were from PDB ID 4C83 (Parker et al. Glycobiology 2014). The structures were superimposed based on Fab backbone atoms using PyMol 3.1 (https://pymol.org). Heavy chains are shown in green and light chains in aqua. Both OSs bind to the respective Fabs primarily through the Fab heavy chain and involve the GlcNAc extension from HepII (γ chain). In the gonococcal OS-Fab complex, the lactose (Glc-Gal) extension from HepII (β chain) plays a major role in binding the Fab.

**Table S1.** X-ray data collection and refinement statistics

| Parameter | Chimeric Fab2C7-OS complex |
|---|---|
| Wavelength | 1.116 |
| Resolution range | 64.07 - 1.6 (1.64 - 1.6) |
| Space group | C 1 2 1 |
| Unit cell | |
| lengths (Å) | 129.52 71.65 116.60 |
| angles (°) | 90.00 114.98 90.00 |
| Total reflections | 1023479 (16975) |
| Unique reflections | 164792 (4461) |
| Multiplicity | 6.2 (3.8) |
| Completeness (%) | 99.92 (99.90) |
| Mean I/sigma(I) | 13.04 (0.9) |
| Wilson B-factor | 28.58 |
| R-merge | 0.05891 (1.686) |
| R-meas | 0.06414 (2.031) |
| R-pim | 0.02499 (1.118) |
| CC1/2 | 1 (0.387) |
| Reflections used in refinement | 127405 (9089) |
| Reflections used for R-free | 1987 (124) |
| R-work | 0.1931 (0.3595) |
| R-free | 0.2104 (0.3529) |
| Number of non-hydrogen atoms | 7376 |
| macromolecules | 6317 |
| ligands | 207 |
| solvent | 852 |
| Protein residues | 833 |
| RMS(bonds) | 0.017 |
| RMS(angles) | 1.07 |
| Ramachandran favored (%) | 96.83 |
| Ramachandran allowed (%) | 3.17 |
| Ramachandran outliers (%) | 0.00 |
| Rotamer outliers (%) | 0.00 |
| Clashscore | 0.24 |
| Average B-factor | 34.4 |
| macromolecules | 33.3 |
| ligands | 45.7 |
| solvent | 39.79 |

[a] Statistics generated using Phenix "Generate Table 1 for journal." Statistics for the highest-resolution shell are shown in parentheses

**Table S2.** $^{1}H$ and $^{13}C$ NMR data of the purified octasaccharide (ppm)

| | H1 | H2 | H3 | H4 | H5 | H6 | H6' | H7/H7' | H8/H8' |
|---|---|---|---|---|---|---|---|---|---|
| **βGal (A)** | 4,39 | 3,51 | 3,66 | 3,88 | 3,72 | 3,93 | 3,70 | / | / |
| | 102,9 | 71,0 | 72,6 | 68,7 | 75,5 | 60,5 | 60,5 | / | / |
| **βGal (B)** | 4,43 | 3,49 | 3,62 | 4,07 | 3,68 | 3,90 | 3,72 | / | / |
| | 103,0 | 71,0 | 72,6 | 65,7 | 75,5 | 60,5 | 60,5 | / | / |
| **αGlc (C)** | 4,49 | 3,39 | 3,58 | 3,48 | 3,51 | 3,76 | 3,68 | / | / |
| | 102,1 | 78,2 | 74,3 | 79,1 | 75,3 | 64,1 | 64,1 | / | / |
| **αGlcNAc (D)** | 5,05 | 3,83 | 3,22 | 3,43 | 3,80 | 4,49 | 4,12 | / | / |
| | 99,4 | 54,3 | 71,0 | 69,8 | 71,6 | 64,0 | 64,0 | / | / |
| **βGlc (E)** | 5,30 | 3,60 | 3,74 | 3,64 | 3,80 | 3,76 | 3,68 | / | / |
| | 100,2 | 71,3 | 71,1 | 77,8 | 71,1 | 61,3 | 61,3 | / | / |
| **αHepI (F)** | 5,82 | 4,09 | 4,17 | 3,64 | 4,00 | 4,07 | / | 3,75/3,69 | / |
| | 97,9 | 78,6 | 74,7 | 71,7 | 71,1 | 71,4 | / | 63,1 | / |
| **αHepII (H)** | 4,97 | 4,07 | 3,97 | 4,01 | 3,64 | ? | / | / | / |
| | 100,6 | 68,2 | 71,8 | 75,8 | 69,1 | ? | / | ? | / |
| **KDO (K)** | / | / | 2,00/1,86 | 4,07 | 4,00 | 4,05 | / | 4,20 | 3,66/3,62 |
| | / | / | 34,3 | 65,7 | 68,3 | 74,6 | / | 66,4 | 62,7 |

**Table S3.** Experimental and calculated proton distances of gonococcal octasaccharide

| Distance | Experimental[a] | MD simulations[b] |
|---|---|---|
| A1-A3 | 2.5 | 2.7 |
| E1-F3 | 2.5 | 2.8 |
| C1-H4 | 2.3 | 2.4 |
| A1-E6 | 2.4 | 2.5 |
| D1-F2 | 2.3 | 2.3 |
| C1-H6 | 2.3 | 2.1 |
| B1-C4 | 2.4 | 2.4 |
| A1-E4 | 1.7 | 2.6 |

[a] Experimental inter-proton distances for OS (NOESY, estimated error 5–10%)
[b] Calculated inter-proton distances for octasaccharide in the bound state (ensemble average from MD simulation)

**Table S4.** Interactions between Fab 2C7 and octasaccharide based on crystal structure

| OS residue | Heptose extension | Fab chain | Number of H bonds[b] | Interaction area ($Å^2$)[b] | Average buried surface area ($Å^2$) |
|---|---|---|---|---|---|
| GMH 3 | NA[c] | heavy | 0 / 0 | 25.6 / 18.4 | 22.0 |
| BGC 4 | alpha | heavy | 0 / 0 | 32.5 / 34.3 | |
| GAL 5 | alpha | heavy | 0 / 0 | 71.2 / 70.3 | 105.3 |
| GAL 5 | alpha | light | 0 / 0 | 0.6 / 1.7 | |
| BGC 6 | beta | heavy | 2 / 2 | 100.2 / 99.0 | |
| GAL 7 | beta | heavy | 2 / 2 | 141.0 / 140.7 | 294.5 |
| GAL 7 | beta | light | 0 / 0 | 54.0 / 54.0 | |
| NDG 8 | gamma | heavy | 4 / 4 | 134.3 / 134.8 | 134.6 |
| **Total** | | | | | 556.4 |

[a] Interactions analyzed with Proteins, Interfaces, Structures and Assemblies (PISA) server (Krissinel and Henrick J Mol Biol 2007)

[b] Values correspond to the two copies in the asymmetric unit (heavy chains A, C; light chains B, D; OS E, F

[c] NA, not applicable

**Table S5.** H bonds between Fab 2C7 heavy chain and octasaccharide based on crystal structure

| **OS residue / atom** | **Fab residue / atom** | **N-O distance (Å)[a]** |
|---|---|---|
| **Beta chain** | | |
| BGC 6 O2 | N76 HD21 | 3.3 / 3.3 |
| BGC 6 O3 | N76 HD22 | 3.1 / 3.1 |
| BGC 6 O3 | N57HD21 | 3.1 / 3.1 |
| GAL 7 O2 | Y126 H (backbone amide) | 3.0 / 3.2 |
| GAL 7 O4 | N57 HD22 | 3.2 / 3/1 |
| **Gamma chain** | | |
| NDG 8 O3 | R124 HE | 3.1 / 2.9 |
| NDG 8 O4 | R124 HH21 | 3.1 / 3.4 |
| NDG 8 O7 | W125 H (backbone amide) | 2.9 / 3.0 |
| NDG 8 O6 | W125 HE1 | 3.2 / 3.2 |

[a] Two values correspond to the two copies in the asymmetric unit (heavy chain: A, C; OS: E, F)

**Table S6.** Water-mediated H bonds between Fab 2C7 and octasaccharide based on crystal structure

| **OS residue / atom** | **$H_2O$ atom**[a] | **Fab residue / atom** | **OS-$H_2O$ distance (Å)**[a] | **$H_2O$-Fab distance (Å)**[a] |
|---|---|---|---|---|
| GMH 3 O4 | 106 / 147 | N76 ND2 | 3.2 / 3.2 | 3.0 / 3.0 |
| GMH 3 O7 | 6113 / 6245 | T54 O | 2.8 / 3.3 | 3.4 / 3.2 |
| GMH 3 O7 | 6063 / 6244 | N78 OD1 | 3.4 / 3.6 | 2.7 / 2.8 |
| GLC 6 O2 | 106 / 147 | N76 ND2 | 3.1 / 3.0 | 3.0 / 3.0 |
| GLC 6 O2 | 124 / 447 | N79 OD2 / ND2 | 2.7 / 2.8 | 3.1 / 3.2 |
| GAL 7 O6 | 6165 / 402 | S83 OG | 2.8 / 2.9 | 3.0 / 2.8 |
| GAL 7 O6 | 609 / 6246 | Y126 OH | 2.9 / 3.0 | 2.8 / 2.7 |
| NDG 8 O4 | 6041 / 6455 | D55 OD1 | 2.7 / 2.9 | 3.1 / 2.8 |
| NDG 8 N2 | 10 / 95 | T54 O | 3.0 / 2.9 | 3.5 / 3.6 |

[a] Two values correspond to the two copies in the asymmetric unit (heavy chain A, C; OS E, F); distance measured between non-H atoms

**Table S7.** Comparison of Fab 2C7 residues interacting with octasaccharide or cyclic peptide 2 (CP2)[a]

| **Fab Residue[b]** | **OS (chains E/F) (PDB ID 9Y5C)** | | | **CP2 (chains F/E) (PBD ID 8DUZ)** | | |
|---|---|---|---|---|---|---|
| **Heavy chain (A/C)** | Buried Surface Area (Å$^2$) | Solvation energy, D$^i$G (kcal/mol) | H-bond | Buried Surface Area (Å$^2$) | Solvation energy, D$^i$G (kcal/mol) | H-bond |
| Asp55 (H1) | 25.0 / 25.6 | -0.12 / -0.15 | No / No | --- / ---[c] | --- / --- | --- / --- |
| Asn57 (H1) | 52.2 / 51.0 | -0.54 / -0.54 | No / Yes | 39.7 / 41.4 | -0.48 / -0.48 | Yes / Yes |
| Asn76 (H2) | 36.8 / 37.6 | -0.39 / -0.40 | Yes / No | 35.4 / 32.9 | -0.28 / -0.31 | Yes / Yes |
| Asn79 (H2) | 4.8 / 6.1 | -0.05 / -0.07 | No / No | 27.3 / 23.1 | -0.30 / -0.26 | Yes / Yes |
| Phe81 (H2) | --- / --- | --- / --- | --- / --- | 102.8 / 86.2 | 1.64 / 1.38 | No / No |
| Arg124 (H3) | 34.1 / 30.2 | 0.05 / 0.06 | Yes / Yes | --- / --- | --- / --- | --- / --- |
| Trp125 (H3) | 182.4 / 181.7 | 1.19 / 1.10 | Yes / Yes | 86.9 / 86.0 | 0.93 / 0.93 | Yes / Yes |
| Tyr126 (H3) | 82.4 / 83.2 | 0.75 / 0.75 | Yes / Yes | 77.3 / 72.8 | 0.51 / 0.51 | No / No |
| **Light chain (B/D)** | | | | | | |
| Trp113 (L3) | 33.6 / 33.8 | 0.54 / 0.54 | No / No | 36.0 / 35.7 | 0.56 / 0.57 | No / No |
| Trp118 (L3) | 23.3 / 23.2 | 0.11 / 0.11 | No / No | --- / --- | --- / --- | --- / --- |

[a] Interaction data obtained from Proteins, Interfaces, Structures and Assemblies (PISA) web server (http://www.ebi.ac.uk/pdbe/pisa)(Krissinel and Henrick, 2007). Replicate values are for two independent copies of heavy chains (A,C) or light chains (B,D).

[b] Fab residues with buried surface area >20.0 Å$^2$ in at least one Fab complex are listed. H1-H3 and L3 refer to the CDR loops of the heavy and light chains, respectively.

[c] ---, no interaction
